# The Cellular Expression and Genetics of an Established Polymorphism in *Poecilia reticulata*; “Purple Body, (*Pb*)” is an Autosomal Dominant Gene

**DOI:** 10.1101/121277

**Authors:** Alan S. Bias, Richard D. Squire

**Affiliations:** Independent Researcher and Swordtail Guppy Breeder. Mailing address: P.O. Box 1508, Lewisburg, West Virginia 24901, USA. http://orcid.org/0000-0002-9093-619X.; Biology Department (retired), University of Puerto Rico, Mayaguez campus, Mayaguez, Puerto Rico, USA. Mailing address: P. O. Box 3227, Mayaguez, P.R., USA 00681-3227. http://orcid.org/0000-0002-3916-0672.

**Keywords:** Guppy color and modifications, chromatophore, violet iridophore, blue iridophore, violet-blue iridophore, xanthophore, xantho-erythrophore, Purple Guppy, Purple Body gene, Metal Gold Iridophore, *Poecilia reticulata*, *Poecilia obscura*, *Poecilia wingei*, Cumana’ Guppy, Campoma Guppy, Endler’s Livebearer.

## Abstract

Modification of wild-type carotenoid orange and pteridine red coloration and spotting of male ornaments in both wild populations of *Poecilia reticulata* (Guppies) and modern Domestic Guppy strains by the Purple Body gene has long been overlooked in research articles and little understood in breeder publications. This modification is commonly found in wild-type *Poecilia reticulata reticulata* populations from numerous collection sites and has been photographed but not recognized in these collections. It is non-existent or near absent in collections taken from variant populations of *Poecilia reticulata wingei*. We identify and determine the mode of inheritance, cellular and phenotypic expression by the Purple gene in these stocks. The Purple Body color pigment modification is a distinct polymorphism in wild *P. reticulata reticulata* populations. Its existence suggests multiple benefits that satisfy female sexual selection preferences, and minimize or reduce potential predation risks. Photographic and microscopic evidence demonstrated that Purple Body is a normal polymorphism in wild and domestic guppies modifying color pigment regions. Purple Body is inherited as an autosomal incompletely dominant trait.

## Introduction

The intent of this paper is multifold; 1. Identify the mode of inheritance of the Purple Body trait. 2. Provide photographic and microscopic exhibits of Purple Body and non-Purple Body for ease in identification. 3. To encourage future study interest at a cellular level of populations in which Purple Body highlights near-UV (Ultra-Violet) reflective qualities. 4. To stimulate molecular level studies of Purple Body and to identify the linkage group (LG) to which it belongs.

A generally accepted definition of polymorphism states: “(1) Polymorphism is the occurrence together in the same habitat of two or more distinct forms of a species in such proportions that the rarest of them cannot be maintained by recurrent mutation. (2) If a genetically controlled form occurs in even a few percent of a population it must have been favored by selection. (3) Polymorphism may either be transient, in which a gene is in process of spreading through a population unopposed, or balanced, in which it is maintained at a fixed level by a balance of selective agencies. (4) Owing to the recurrent nature of mutation, transient polymorphism is generally due to changes in the environment, which make the effects of a previously disadvantageous gene beneficial (Ford 1945).”

Wild *Poecilia reticulata*, both in native populations and feral introductions, exist in a previously undocumented polymorphic state; Purple Body and non-Purple Body. In Domestic strains both polymorphisms persist as a direct result of intended breeder intervention and as an unintended result of outcrosses between fixed phenotypic strains. Therefore, it is safe to assume that the co-existence of two sympatric phenotypes, in both wild and domestic stocks, must confer a selective advantage. The two most likely possibilities are reduction in predation and/or favoritism in sexual selection.

For nearly 100 years published research has focused on “heritability, brightness, intensity of orange chroma and hue in male ornaments (Winge 1922a, 1927; Lindholm 2002; Deere 2012), most often in the context of sexual selective preference or environmental carotenoid intake. Over the last decades we have seen a gradual shift towards identification of benefits derived from the reflective qualities of iridophore biased phenotypes. This frequently involved the study of Opsin, and UV and near-UV spectrum vision in Guppies and their predators. Additional consideration was given to the nature of ornaments being composite traits with independent origins (Grether 2004).

*P. reticulata* pteridine based color appears as shades of red, while carotenoid color is more yellow-orange. The "wild-type" guppy red is maintained in a variety of environments by synthesizing sufficient pteridines to balance the amount of dietary carotenoids obtained from food sources. Iridophore development begins in embryonic stages, continues with onset of sexual maturity and is completed in young adulthood. Reflective qualities of iridophores increase with maturation and development of guanine crystals (Gundersen 1982; Deere 2012). Intensity and reflective qualities of coloration are complemented by both underlying basal and ectopic melanophores.

Studies on extreme polymorphic variation found within Poeciliid species over the last 100 years abound involving *P. reticulata, P. parae, P. picta*. For the most part those within wild *Poecilia reticulata* and *Poecilia reticulata wingei* have focused on “wild-type” orange color pigmentation of spotting; its benefits from carotenoid intake, Y-linked inheritance for spotting, female sexual selection preferences, and heterogametic benefits in the form of reduced predation on females. Of late emphasis has seen a shift towards benefits derived from both UV and near-UV coloration.

Xantho-erythrophores primarily utilize the absorbed carotenoids ingested from local dietary resources. Carotenoid availability produces variability in expression within and between populations (Grether 1999). This is in contrast to red drosopterin based color pigment, which is synthesized by the individual *de novo* (Goodwin 1986). Male orange ornaments are the product of both orange carotenoids and red pteridine color pigments: drosopterins.

A direct correlation between carotenoid intake and higher levels of drosopterin synthesis has been identified. While pteridine red and tunaxanthin carotenoid orange were identified in male spotting, only the carotenoid orange was obtained from the algae in the environment in wild study populations (Grether 2001). Counter-gradient variability found in male orange ornaments from multiple locales has been documented (Grether 2005a, 2005b; Deere 2012) as the result genetic differences between populations. The ratio of carotenoids and drosopterins had a direct effect on size and hue of orange spotting.

Research utilizing Domestic Guppies has proven both X and Y-linked inheritance for red color pigmentation in finnage (Khoo and Phang 1999). Documented results by Domestic Guppy Breeders reveal many strains have been produced with both X and Y-linked inheritance for red color pigment in both body and finnage. Many females capable of passing red in domestic Guppy stocks often express coloration, while others resembling wild color / tail neutral guppy females do not.

Wild-type female guppies in native locales, and domestic strains, lacking coloration are also capable of passing on unexpressed red color pigment to daughters and expressed color pigment to sons (Gordon 2012; Bias, *unpublished breeding results*). Such sexual dimorphism reduces predation on females, while allowing it to varying extents on males. Continued breeding’s in Domestic strains are now showing that red color pigment is not simply the result of an XY-linked mode of inheritance, but are also suggestive of autosomal linkage as well. A recent publication involving red color morphs found within *Poecilia picta* supports this assertion (Lindholm 2015).

The purple phenotype has been present in hobbyist stocks for decades, but has been largely unrecognized except in the case of pure-bred all-purple strains, such as the Roebuck IFGA (International Fancy Guppy Association) Purple Delta described in Materials. The Purple tail guppy with purple iridescence on body and causal fins seems to be the same as the “all-purple” term we use here, and was studied in Ben et al. (2003), where they extensively studied and discussed the enzymatic processes involved in producing various pteridine pigments in the guppy. No formal description of genotype, in regard to the Purple tail guppy, was proffered or elucidated by the authors. Their “preliminary study concentrated on PTPS (*6-pyruvoyl tetrahydropterin synthase*), which is the main rate-limiting enzyme in the pteridine biosynthesis pathway …and XDH (*xanthine dehydrogenase*), which is largely responsible for the catabolism of 7,8-dihydropterin to pigmented pteridines, including isoxanthopterin, xanthopterin, and leucopterin…”

They found that “Purple Tail, Diamond and Blue Metallic have silvery iridescent caudal fins resulting mainly from the presence of iridophores. Purple Tail looks more yellowish than Diamond and Blue Metallic, while Blue Metallic has deeper blue iridescence than Diamond. The PTPS mRNA levels were moderately expressed in these guppy varieties and appeared to be related to the density of the xanthophores.” Purple tail and Diamond had a higher density of iridophores and a higher XDH expression then the other mentioned strains. They finally concluded that the Purple tail phenotype is produced by the action of iridophores that give a reflective blue effect (structural color) in this case, and xanthophores which contain yellow (and sometimes red/orange) pigments in particular.

Ben et al. also discussed the pteridine pigments in other color varieties of the guppy. But they did not discuss purple coloration found in strains with more complex purple, red, orange and yellow colors present in the same fish, nor did they investigate the inheritance of the purple gene. This senior author has presented the differences between “Purple Body”, all-purple, red, and orange color patterns too many hobbyist groups both in the United States and internationally for a number of years. He discovered the microscopic differences between these two traits in chromatophore arrangements associated with red, orange, yellow, and purple spots. After discussions with the junior author, he refined these conclusions and then embarked upon a series of crosses designed to determine how the Purple Body gene is inherited. The junior author conducted the genetic analyses based upon the cross results, and co-authored the final paper.

Breeding tests involving this modification of orange spotting reveal this trait to have an incompletely dominant mode of inheritance. As such a formal name and nomenclature of **Purple Body (*Pb*)** is therefore suggested. *[**Note**: Hereafter Purple Gene and Purple Body Gene are used interchangeably]*.

## Description (Spectral Expression of Pb vs. non-Pb)

A frequent and previously undescribed modification of “wild-type” orange color pigment exists within both wild and domestic stocks of Guppies. Visually, coloration is modified from a highly reflective orange to a “purplish-pink” coloration in Grey (*g*), Blond (*b*) and Golden (*g*) (Goodrich 1944), European Blau (*r*) (Dzwillo 1959) and Asian Blau (*Ab*) (*Undescribed* - see Bias 2015) variants, most vividly noticeable in grey and blond.

While this purplish-pink modification of individual and pattern spotting is readily visible in color drawings and photo plates of prior published results, descriptions have been generally limited to gene(s) at specific loci in reference to iridescence, red, orange, purpureus and rubra. Thus, indicating no formal documentation for the existence of a modifier gene of orange spotting (*see more later*). To further rule out the possibility of prior formal Pb description, an exhaustive review was made of published research to date. This including any early descriptions closely associated to “purple” and “red” coloration in body and finnage. Those identified include:

> A. Purple Tail; The phenotypic name for an Asian commercial strain referenced in the 6-pyruvoyl tetrahydropterin synthase (*PTPS*) mRNA study. The pigment phenotype was simply listed as “Purple iridescence on body and caudal fins.” No formal description of trait(s) was made (Ben 2003).
> B. Purpureus (*Pu*); Yellow and red dorsal pattern description (Natali and Natali 1931; Kirpichnikov 1981). No reference was made to modification of red color pigment.
> C. Rubra (*ru* or *r*); 1. Red color proximally in the upper edge of the caudal fin. 2. Large oblong red side spot, lying for the most part below and behind the dorsal fin. 3. Dark side-dot in the tail at the base of the caudal fin (Winge 1922b). No reference made to modification of red color pigment.
> D. Anterior Rubra (*ar*); Y-link mode of inheritance and allelic to ru complex (Blacher 1928). No reference made to modification of red color pigment.

High resolution photography and microscopic study shows the co-existence of varying populations of both violet and blue structural iridophores in all individuals, both male and female (*Bias and Squire 2017b, Pb Microscopy Study, forthcoming*). Progressive focal adjustment failed to alter this violet coloration to blue. This is consistent with current Guppy research (*Figure 6C*, Kottler 2014; *Figure 5A’*, Kottler 2015) Betta research (*Figure 1b* Khoo and Phang 2013), and Zebrafish research (Figure 3, Patterson 2013; Figure 1H, Mahalwar 2014; Figure 2 and 3, Singh 2015). Violet and blue structural iridophores and melanophores are always found in close proximity with one another. Forming a type of chromatophore unit [***Note:*** *hereafter referenced as violet-blue (iridophores) for ease of discussion*]. Violet-blue iridophores **(Fig 2)** are most visible along the topline and in between regions lacking a clearly defined silver iridophore pattern, often including caudal-peduncle base.

**Fig 1.**
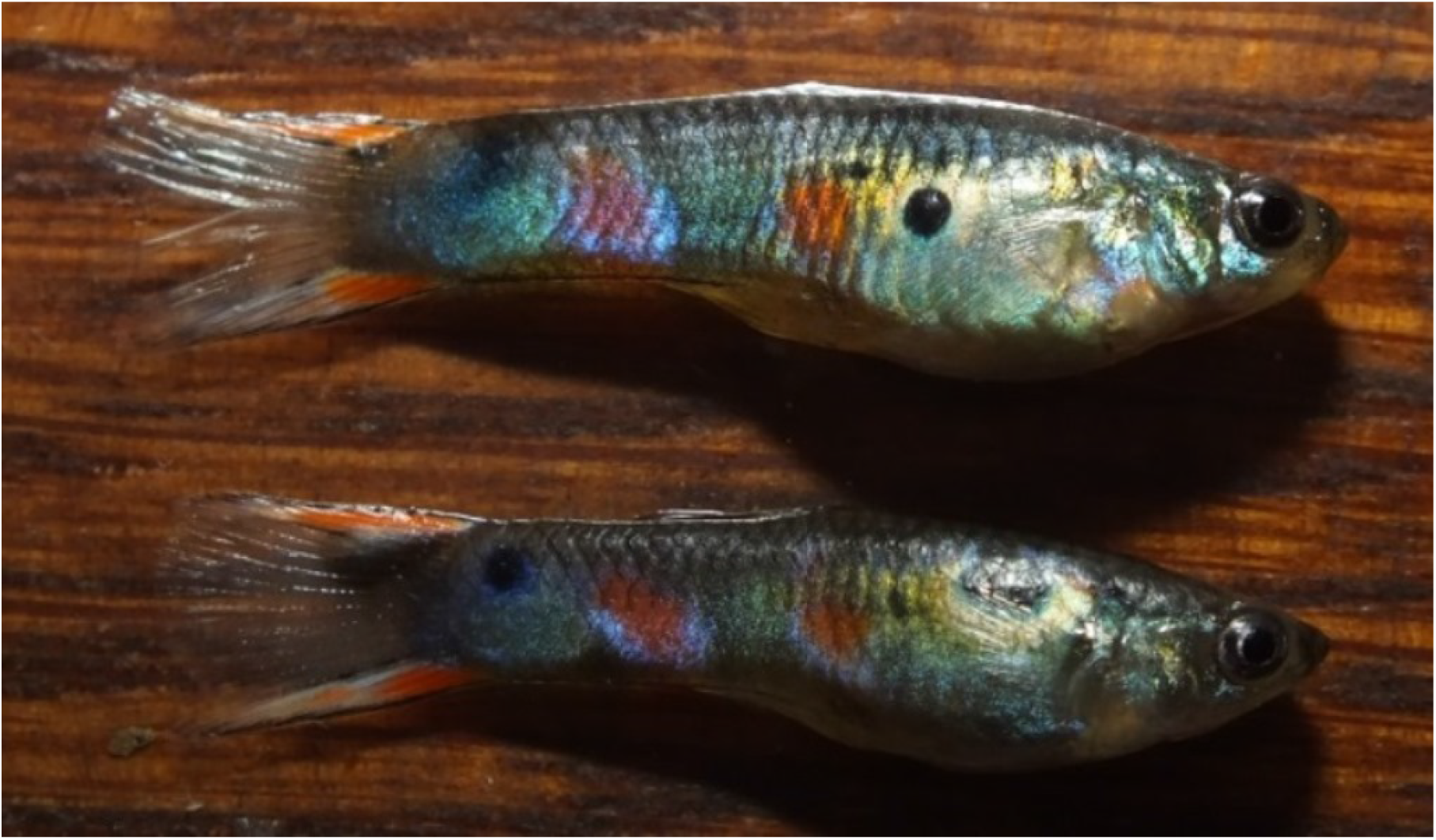
Wild-type Purple Body (*Pb*) males in heterozygous condition.

**Fig 2.**
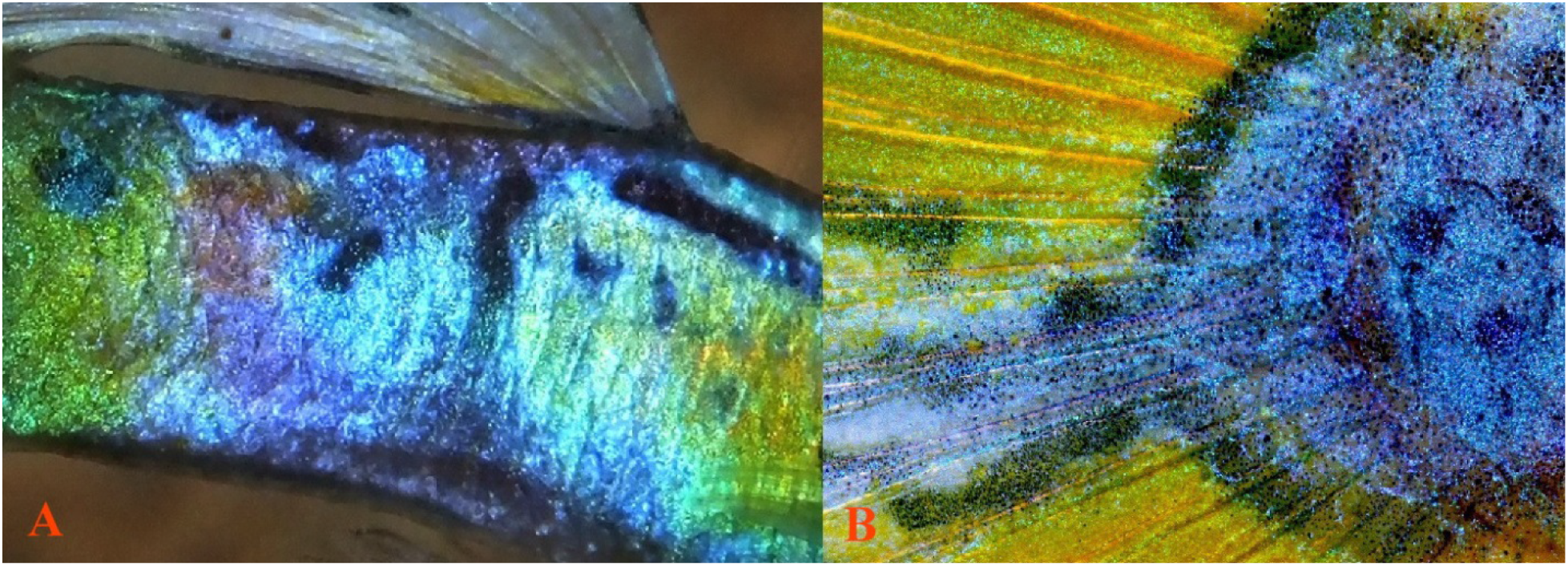
**(A)** Violet-blue Iridophores in peduncle, **(B)** Violet-blue iridophores in caudal, photo courtesy of Christian Lukhaup (right).

**Fig 3.**
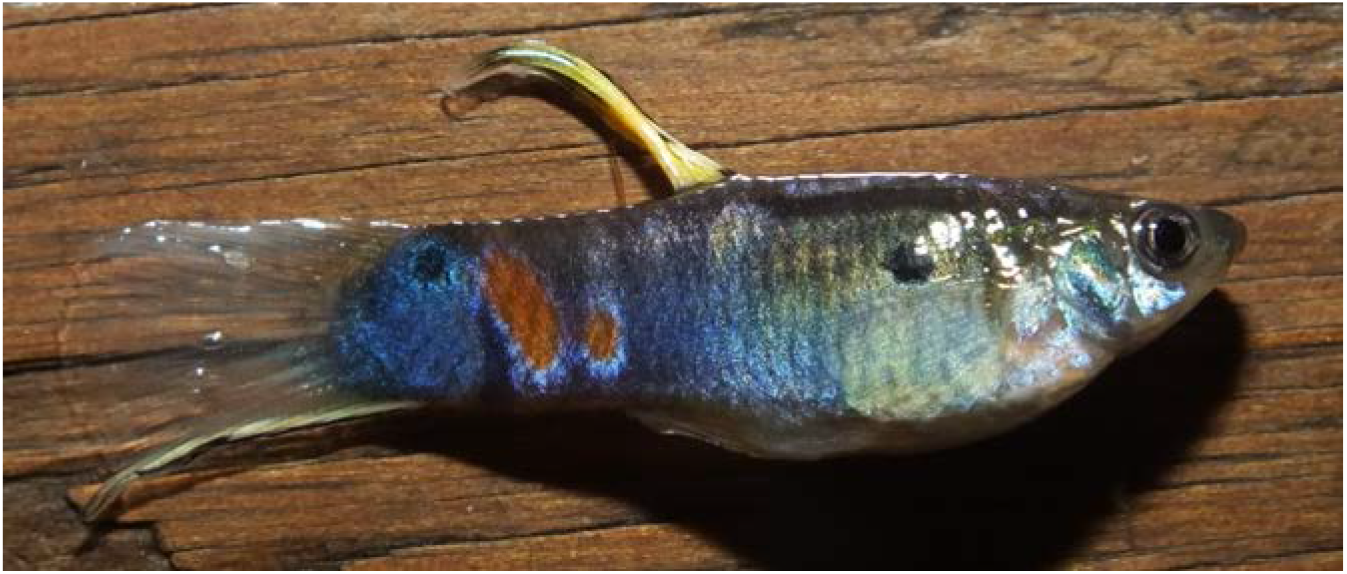
Homozygous Pb.

**Fig 4.**
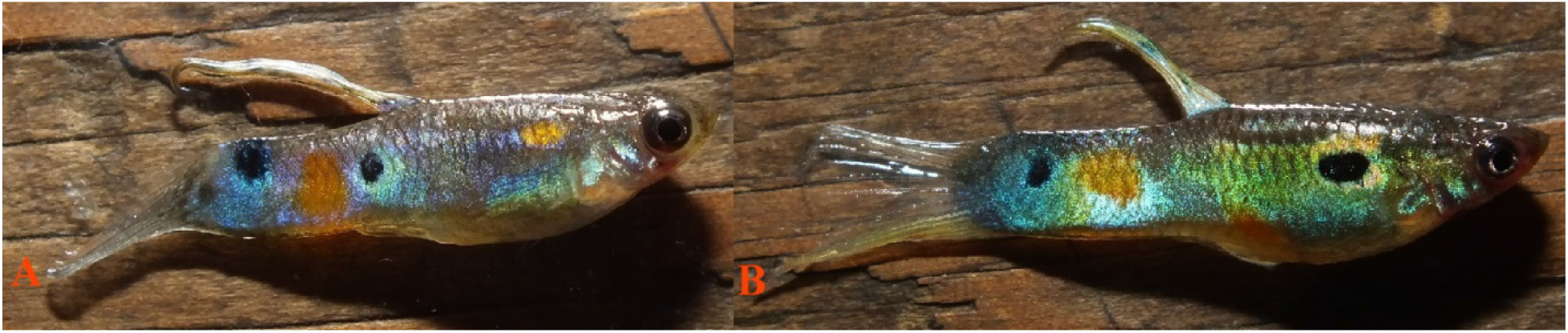
**(A)** Heterozygous Pb (left), **(B)** Non-Pb male (right).

**Fig 5.**
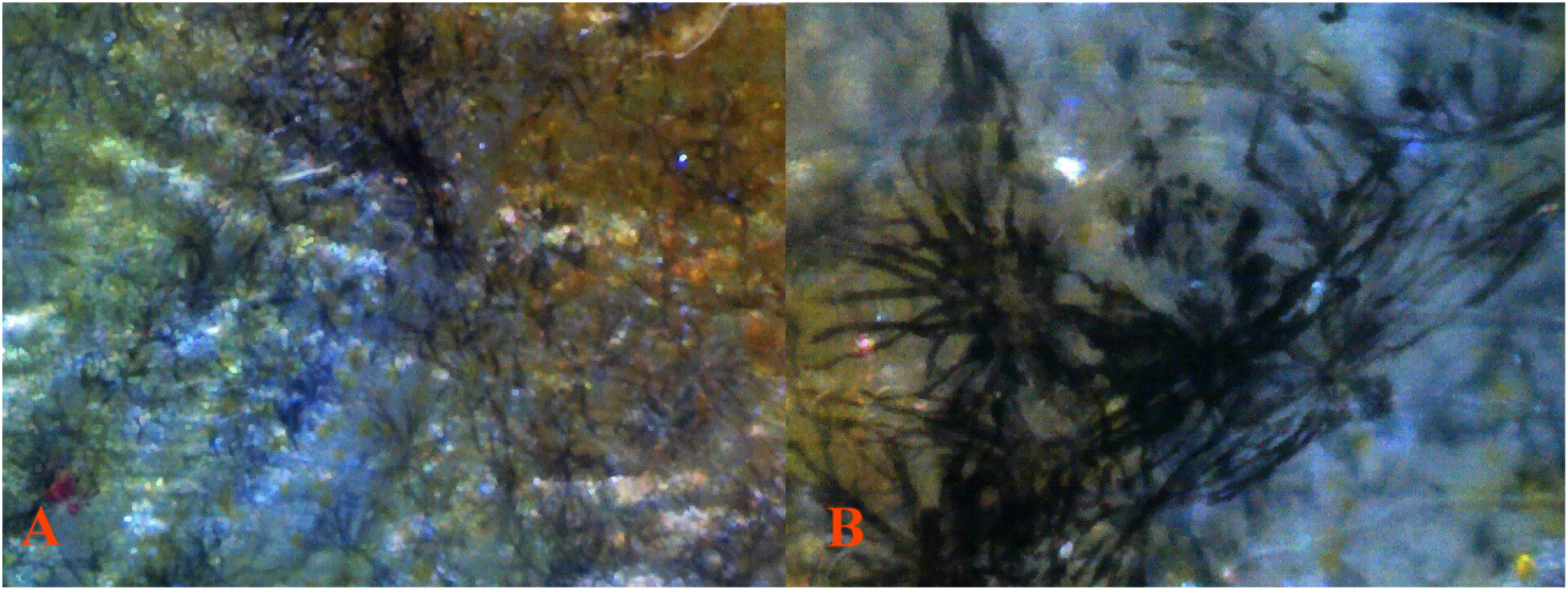
**(A)** 8 Pb 40× 11 under reflected light without transmitted light *(dissected)*. **(B)** 8 Pb 100× 12 under reflected light without transmitted light *(dissected)*. Pb modified dendritic melanophore strings and violet-blue iridophore chromatophore units along scale edging producing reticulated pattern. The extreme “length” of dendrites is a result of Pb.

**Fig 6.**
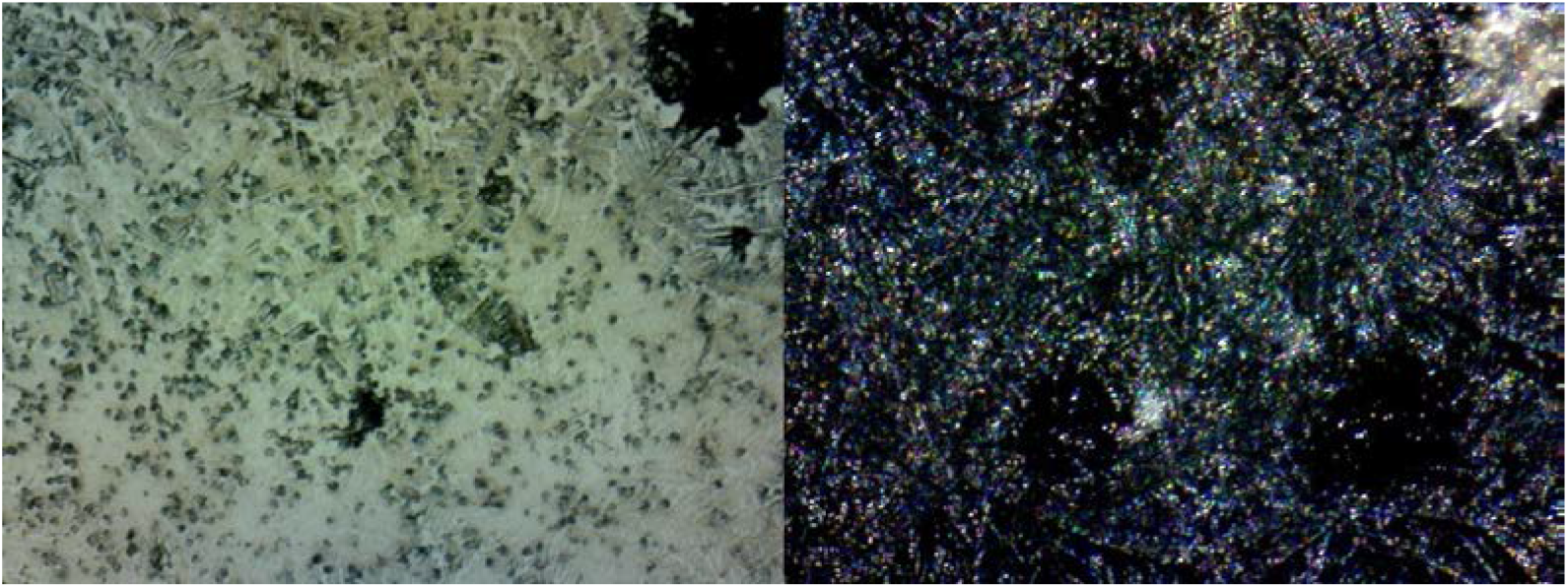
Wet mounts, no cover glass, dehydrated. Aqueous humor fluid extraction **(A)** 32 40 8 *Pb/-(dissection)* transmitted light. **(B)** The same field reflected light with white balance adjusted.

Pb modification, zygosity dependent, removes certain classes of yellow-orange-red color pigment over silver iridophores or white leucophores. The term we use here, “white leucophore”, refers to a pigment cell type that was extensively studied both structurally and biochemically in Takeuchi (1976) and identified as a fourth chromatophore type. However Kottler, et al (2014) were not able to confirm the presence of leucophores based upon their physical structure. We were not able to determine whether the cells we observed were white leucophores or white iridophores, but will continue to refer to them here as leucophores. Pb modifies “other existing” color in both body and fins, thus suggestive of being a “full body” modifier, in homozygous fashion. Dark red pteridine color pigment does not seem to be modified by Purple Body in fins lacking an underlying silver iridophore or white leucophore pattern. Modification by Pb seems limited predominantly to wild-type orange color pigment; i.e. that which also contains yellow carotenoids in addition to red pteridines, over an iridophore pattern.

Pb is always found in all-purple fish, but is not by itself sufficient to produce the all-purple phenotype in heterozygous expression. Homozygous Pb expression resulting in further removal of xantho-erythrophores, in conjunction with both increased populations and/or the visibility of modified melanophores and naturally occurring violet-blue iridophores, is required for production of the all-purple phenotype. Violet is a true wavelength color, while Purple is a composite produced by combining blue and red wavelength colors. Thus, the violet-blue chromatophore unit and removal of xanthophores is required to produce an all-purple phenotype.

Based on breeding results, phenotypic observation and microscopic study, further observations are offered. Expressivity of Pb is not only highly variable, but also zygosity dependent. Yet, the frequency of Pb heterozygotes in wild populations appears high. Zygosity dependent (*Pb/pb or Pb/Pb*), and specific to individual zones of regulation with variability; Pb causes a large reduction on yellow color pigment populations (*xanthophores*). It thus produces a modified purplish-pink expression instead of the orange color pigment (*xantho-erythrophores*). By nature, yellow color pigment in Guppies is highly motile and mood dependent while red color pigment is non-motile. While red color pigment (*erythrophores*) is not altered, or at least altered to a lesser degree, a corresponding noticeable increase in the visibility (possibly increased population levels) of structural violet-blue iridophores is evident, resulting in the increased reflective qualities of individuals.

When not masked by additional color and/or pattern traits, the identification of Purple Body (*Pb*) in both wild-type and domestic males can be easily accomplished through visual phenotypic observation. In non-Purple Body (*pb/pb*) the individual's carotenoid orange color pigment can be described as being vivid bright orange, structurally comprised of densely packed yellow and orange xantho-erythrophores, normally extending to the very edge of the spot. Though coverage over additional iridophore patterns may appear incomplete.

Heterozygous Purple Body (*Pb/pb*) males express modified purplish-pink spots under early morning and late afternoon ambient sunlight. This same coloration can be easily observed under hand held incandescent lighting focused toward the side of the tank. During periods of midday sunlight or under overhead artificial light sources Pb may appear as dark or faded orange.

A reduction in the number of yellow xanthophores results in a corresponding reduction in overall size of individual spotting ornaments. This reveals a “circular ring” produced by an underlying iridophore layer. This well-defined layer of iridophores is the actual underlying precursor required for definition of shape over which color pigment cells populate. In general, spectral observation and photography of guppies from crosses reveal iridophore migration and pattern formation prior to the formation of xantho-erythrophores. Microscopy reveals minimal xantho-erythrophore populations to be already in place at first (*Bias and Squire, 2017b, Pb Microscopy Study forthcoming*). It has been suggested that, “it is crucial to consider iridophores as well, which might attract melanophores and xanthophores to the locations where spots arise during male color pattern formation. Depending on the location, iridophores might also repulse xanthophores or melanophores, or influence their survival” (Kottler 2014). Our findings concur with this suggestion. As will be further discussed in the results, Purple Body has shown an incompletely dominant mode of inheritance. In heterozygous condition (*Pb/pb*) a distinct result is generated while in homozygous condition (*Pb/Pb*) these results are further amplified.

Heterozygous Pb in both wild-type and domestic individuals alters orange spots in select regions of the body and in finnage to “purplish-pink”. Thus, it may not act as a “full body” modifier in heterozygous form. Heterozygous Pb does not appear to greatly reduce visible structural yellow color pigment cells over white leucophore or reflective clustered yellow cells, known in breeder circles as Metal Gold (*Mg*) (*Undescribed* - see Bias 2015), in body and finnage. A slight increase in visibility of violet-blue iridophores is often detected. Additionally noted is an increase and modification in existing melanophore structure and populations.

Homozygous Pb in both wild-type and domestic individuals alters all orange spots found in the body and in finnage to “purplish-pink”. Thus, Pb should be considered a “full body” modifier. Homozygous Pb produces a purple guppy phenotype. Homozygous Pb removes all visible yellow color pigment over white leucophore, but not Mg in body and finnage. This in turn, produces a dramatic increase in the visibility of wild-type violet-blue iridophores. A dramatic increase and modification in existing melanophore structure and populations is noted. Provided are examples **(Fig 3-4)** of Phenotypic distinction between Pb and non-Pb in wild Guppy populations. For examples in Domestic strains see *Bias and Squire 2017c, forthcoming*.

## Description and Characteristics: cellular expression of Pb vs. non-Pb

In general, while there are microscopic differences, our findings of visual distinctions between Pb and non-Pb are often more consistent, as opposed to microscopic distinctions. Much of this is likely the result of variability in both zygosity and ornament composition between individuals, both among and between populations and strains. Microscopically, structural differentiation between xantho-erythrophores appears minimal, with differences in population levels and collection or clustering of xanthophores. Heterozygous Pb exhibits partial reduction in collected xanthophores, and homozygous Pb exhibits near complete removal of collected and clustered xanthophores. Though, it is noted yellow color cell populations consisting of isolated “wild-type” single cell xanthophores remain intact.

Dendritic melanophores are present in both Pb and non-Pb at various locations in the body. Observation of Pb in heterozygous and homozygous condition, for mature individuals, reveals that ectopic melanophore dendrites are often extremely extended **(Fig 5)**. This occurs either as the result of direct modification by Pb, or indirectly through interactions as a result of xanthophore reductions or removal (Kottler, 2015). Overall dendritic melanophore structure is of a much “finer” appearance as compared to non-Pb. Modified melanophores are more often linked together in “chain-like” strings **(Fig 5)**, as compared to non-Pb, both within and outside of areas defined as reticulation along scale edges. While this study did not directly seek to identify an increase in melanophore populations, it was assumed higher numbers of melanophore structures would be present in Pb. While this may be the case, “darker” appearance in Pb vs. non-Pb appears to largely be the result of modification of existing melanophore structures (corolla and punctate) into extended dendrites. Thus, the number of melanophore structures does not appear to drastically increase, in any given individual, only the size of the structures themselves.

The motile nature of melanosomes in ectopic melanophores may allow for changes in reflective qualities or hue of individuals. In conjunction with zygosity dependent removal of xantho-erythrophores, this may satisfy both female sexual selective preferences for conspicuous pattern of “bright orange” under specific lighting, and maintain crypsis in others (Endler 1978). While frequent evidence of dendritic and/or motile yellow color pigment (xanthophore) structures was detected in this study, none was found for dendritic and/or motile iridophores, outside of violet and blue [*hereafter violet-blue*] iridophore clustering associated with ectopic dendritic melanophores. Violet-blue iridophores are more visible in Pb vs. non-Pb, with variability between populations and strains, based on overall genotype. Specific to varying genotypes of individuals, there appears to be an actual increase in the ratio of violet to blue iridophores, as would be expected in an “all purple phenotype”. Whether there is an actual increase in iridophore population numbers, or simply increase visibility, due to reductions or removal of xanthophores and/or altered melanophores was not addressed in this study. Nor was the issue of the modification of the angles at which crystalline platelets reside beneath iridophore layers and basal level melanophores.

## Cellular Comparison: Ocular Media Pb and non-Pb

Our study revealed that all major classes of chromatophores (melanophores, xanthophores, erythrophores, and violet-blue iridophores) and crystalline platelets were present in the cornea, aqueous humor, vitreous humor, outer lens membrane and possibly the lens itself of *Poecilia reticulat*a. Contrary to visual observations and conventional transmitted light microscopy the cornea, aqueous and vitreous humor are not clear under conventional reflected light microscopy. Each possesses independent populations of static and/or free-floating chromatophores. To the authors’ knowledge this is the first time this has been reported in *P. reticulata*.

We postulate that dense layers of violet-blue iridophores in conjunction with melanophores and xanthophores residing within the cornea-aqueous humor (**Fig 6**)-iris-vitreous humor and the surrounding capsule at the anterior pole of the crystalline lens act as “ocular media filters”, with individuals deriving benefit in the UV and/or near-UV spectrum. Pb will provide benefit at lower wavelengths with increased levels of violet iridophores, and non-Pb will have reduced benefit at slightly higher average wavelengths with balanced violet-blue iridophores. In turn, xanthophores counter balance and provide benefit in the higher wavelengths. [**Note:** For complete ocular microscopy study, see Bias and Squire 2017c, *forthcoming*.]

## Cellular Comparison: melanophore modifications in Pb *Pb/pb* vs. non-Pb *pb/pb* under reflected and transmitted light

Dendritic melanophores in heterozygous and homozygous Pb condition, in mature individuals, reveal that dendritic arm structure is extremely extended and finer in appearance **(Fig 7)**. Dendrites are linked together in “chain-like” strings intermingled with violet-blue iridophores in chromatophores units. Within the rear peduncle spot and surrounding edges a noticeable absence of corolla and punctate melanophores was often evident. This absence was abated in other regions **(Fig 8)** of the body or specific to individuals. The angle of incident lighting and spectral capabilities can alter visual perception, so too can the direction of light. Examples are here presented under both reflected and transmitted lighting, to reveal chromatophore visibility and expression for each.

**Fig 7.**
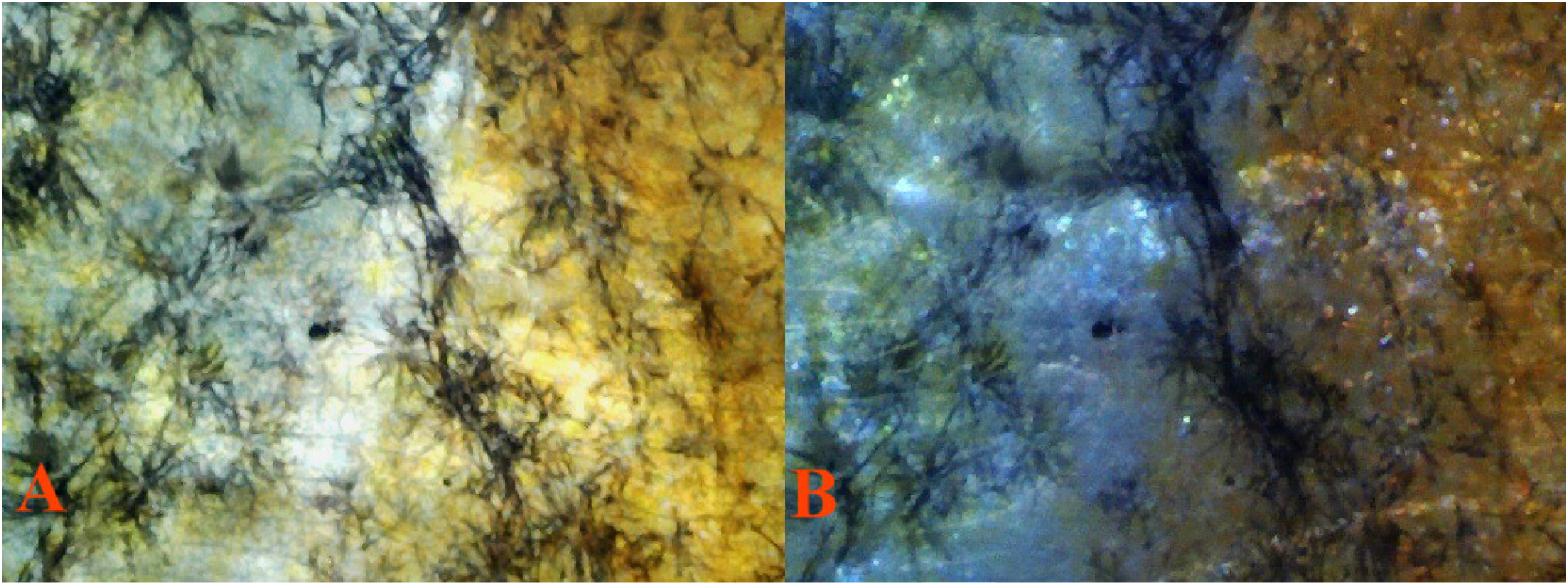
**(A)** 8 Pb 40× 13 *Pb/pb (dissected)* transmitted light. **(B)** The same field, reflected light. Extreme dendritic melanophore modification in reticulated pattern do to heterozygous Pb.

**Fig 8.**
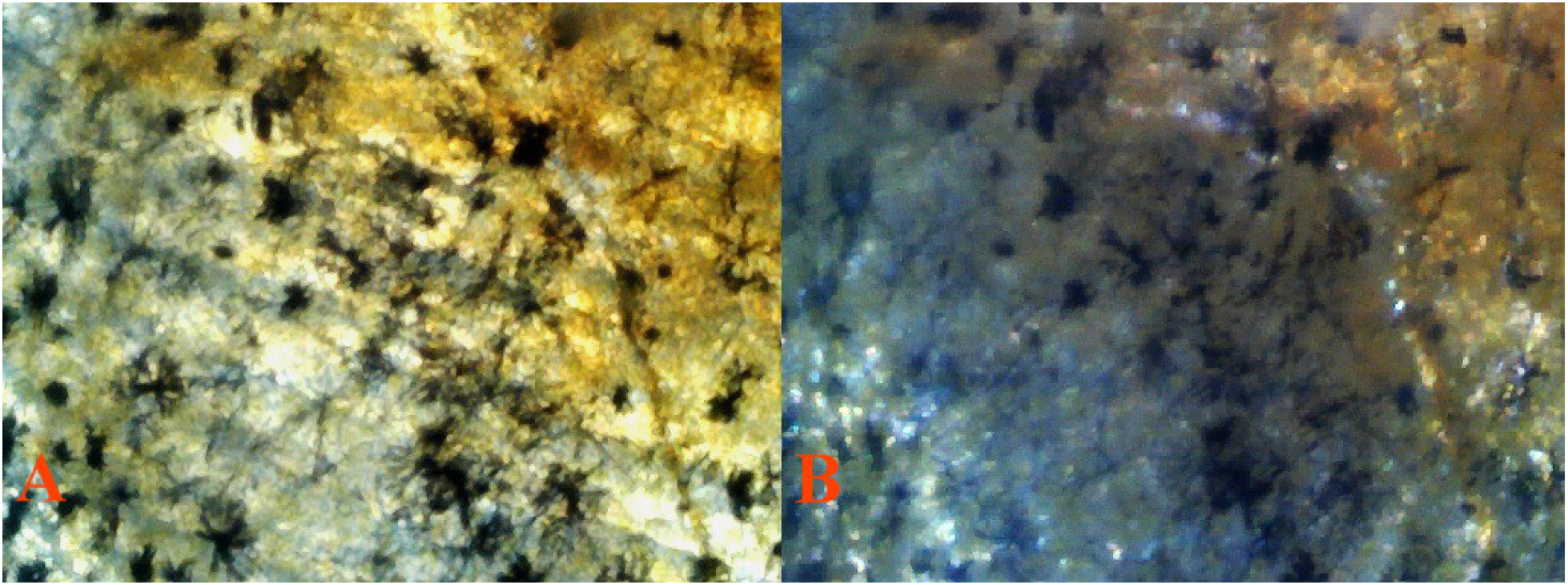
**(A)** 7 non-Pb 40× 15 *pb/pb (dissected)* transmitted light. **(B)** The same field, reflected light. Reduced dendritic melaophore extensions in non-Pb.

## Cellular Comparison: early coloration in Pb *Pb/pb* vs. non-Pb *pb/pb* male Guppies (*Poecilia reticulata*)

The following examples of early coloration in non-Pb and Pb show male expression of violet-blue iridophores macroscopically **(Fig 9)** 100x. Visual distinction is easily made between the two iridophore types. Changes in magnification, progressive focal shift, adjustment in angle of incident lighting or direction of light (reflected or transmitted) consistently failed to remove this visible distinction between violet and blue. Thus, this demonstrates two distinct iridophore populations in the blue-violet spectrum.

**Fig 9.**
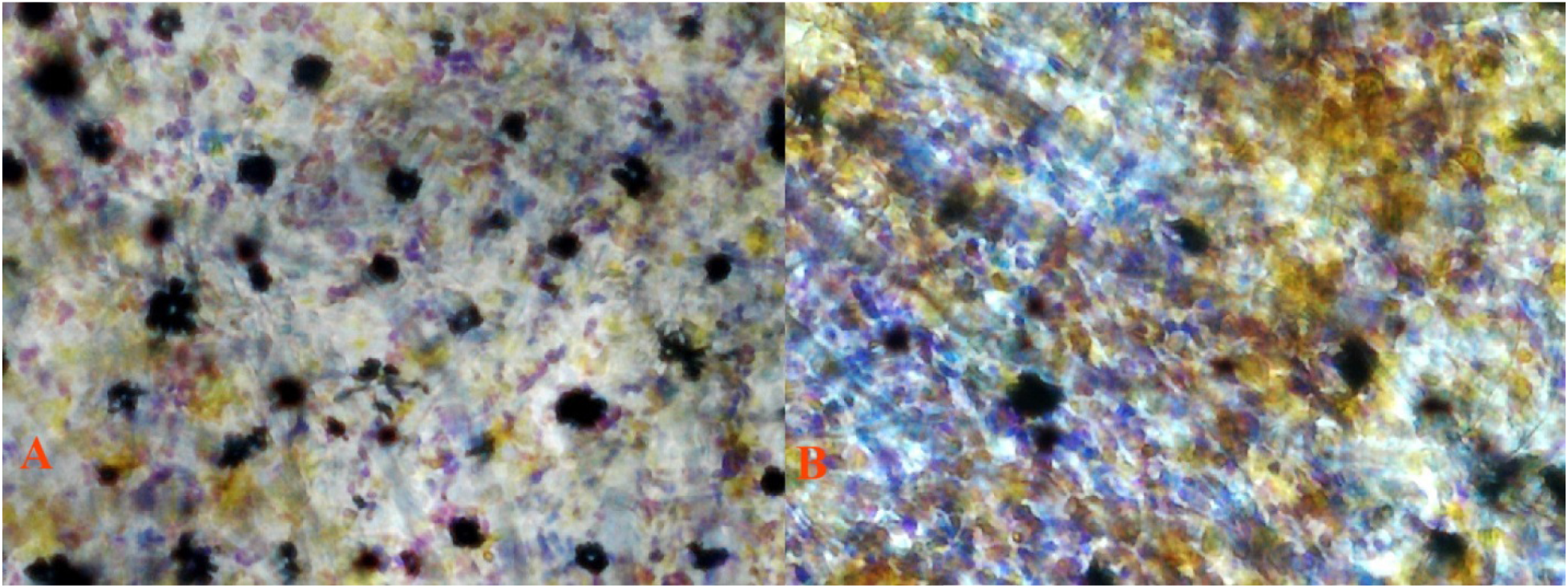
**(A)** 6 Pb 100× 12 *Pb/pb (dissected)* transmitted light. More violet iridophores than blue irdophores. **(B)** 3 non-Pb 100× 14 *pb/pb (dissected)* transmitted light. More evenly distributed blend of violet and blue irdophores in non-Pb as compared to Pb.

Two distinct observations are offered based on early coloration. First, violet and blue iridophores appear “randomly collected among themselves” in similar fashion **(Fig 9)**, as opposed to later mature coloration in which violet and blue iridophores are arranged together in “joined alternating color” groupings in dissimilar fashion. This shows that coloration is nearly complete, while migration to their final location is not. Second, melanophore shape is predominately corolla or punctate in early coloration **(Fig 9)**, as opposed to mature coloration in which dendrites dominate. This indicates that members of the melanophore population are in place, while their final shape is not established. Side by side presentation of similar locations in Pb and non-Pb are presented.

## Cellular Comparison: late coloration in Pb *Pb/pb* vs. non-Pb *pb/pb* male Guppies (*Poecilia reticulata*)

Macroscopically **(Fig 10)** and microscopically **(Fig 11)** visible in heterozygous Pb are partial reductions in collected xanthophores, and in homozygous Pb near complete removal of collected and clustered xanthophores. Yellow color cell populations consisting of isolated “wild-type” single cell xanthophores remain intact.

**Fig 10.**
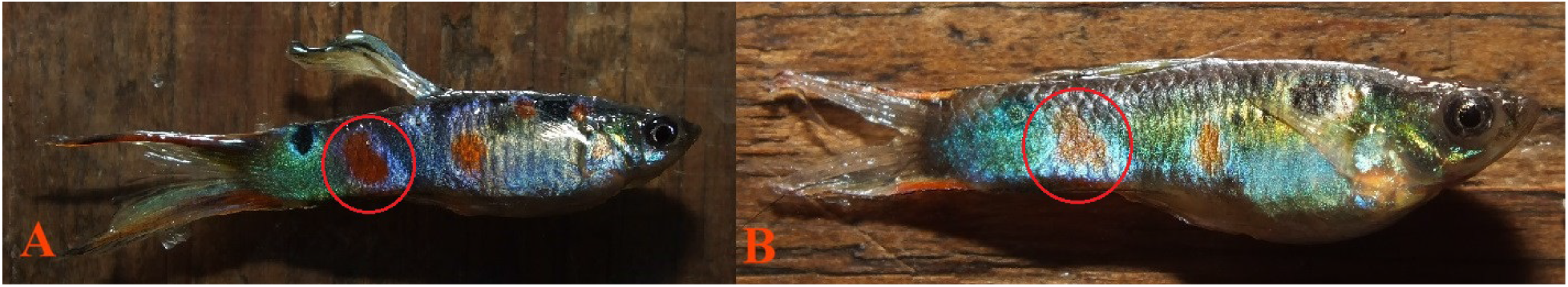
**(A)** Heterozygous 11 Pb male *Pb/pb*, **(B)** 6 non-Pb male *pb/pb*. All slide images taken from posterior orange spot (red circle).

**Fig 11.**
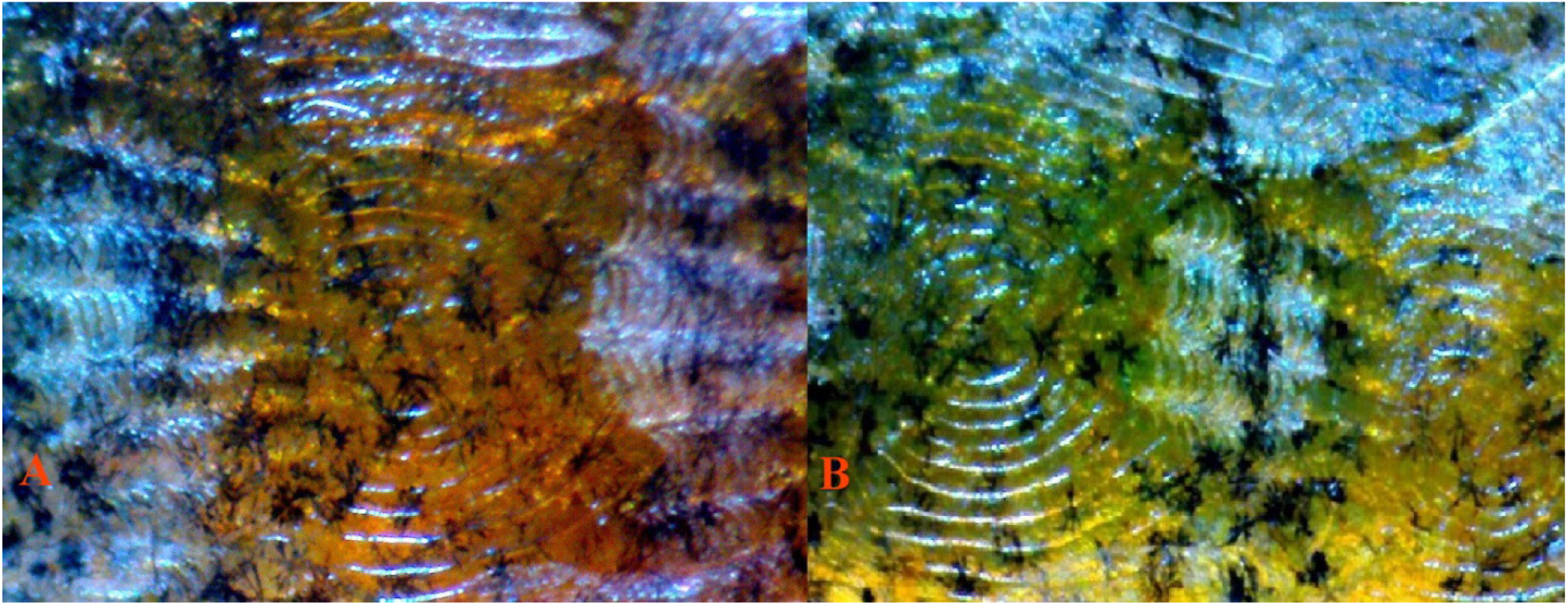
**(A)** 13 Pb 40X 4 *Pb/Pb (dissected)* reflected light. Expected higher visiblity of erythrophores with reduction of xanthophores by Pb modification. **(B)** 14 non-Pb 40X 3 *pb/pb (dissected)* reflected light. Expected evenly distributed xantho-erythrophores in non-Pb.

All major classes of chromatophores were present in the rear peduncle spot and adjoining areas in both Pb and non-Pb. Violet-blue iridophores are more visible in Pb **(Fig 10A)** vs. non-Pb **(Fig10B)**, with variability between study specimens. An increase in the ratio of violet to blue iridophores was observed. Collected and clustered xanthophore populations, found in non-Pb **(Fig 11B)** members of the contemporary group, were reduced in heterozygous Pb **(Fig 11A)** condition and removed in homozygous Pb condition. The retention of isolated xanthophores remained intact in both heterozygous and homozygous Pb condition.

## Cellular Comparison: late coloration in Asian Blau *Ab/ab Pb/Pb* vs. non-Pb *Ab/ab pb/pb* male Guppies (*Poecilia reticulata*)

Asian Blau (*Ab* – *undescribed*, see Bias 2015) presents a unique opportunity to further confirm the spectral removal of yellow color pigment by Pb, though microscopic study reveals this removal to be far from complete. Autosomal incompletely dominant Ab, as opposed to autosomal recessive European Blau (*r* or *r1*, Dzwillo 1959) in heterozygous and homozygous condition removes red color pigment (**Fig 12-14**).

**Fig 12.**
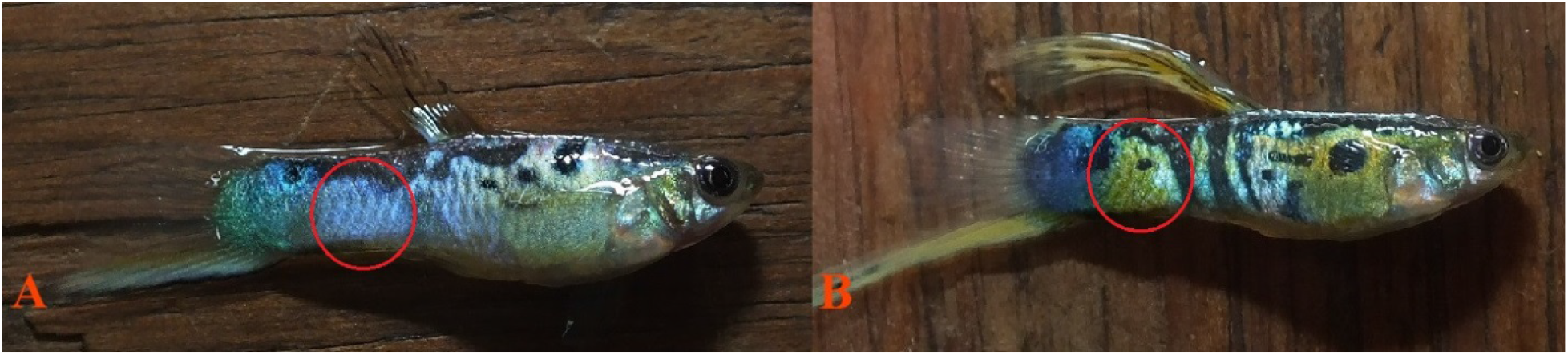
**(A)** 5 Pb male (grey) *Pb/Pb Ab/ab*, **(B)** 4 non-Pb male (grey) *pb/pb Ab/ab*. All slide images taken from posterior modified orange spot (red circle).

**Fig 13.**
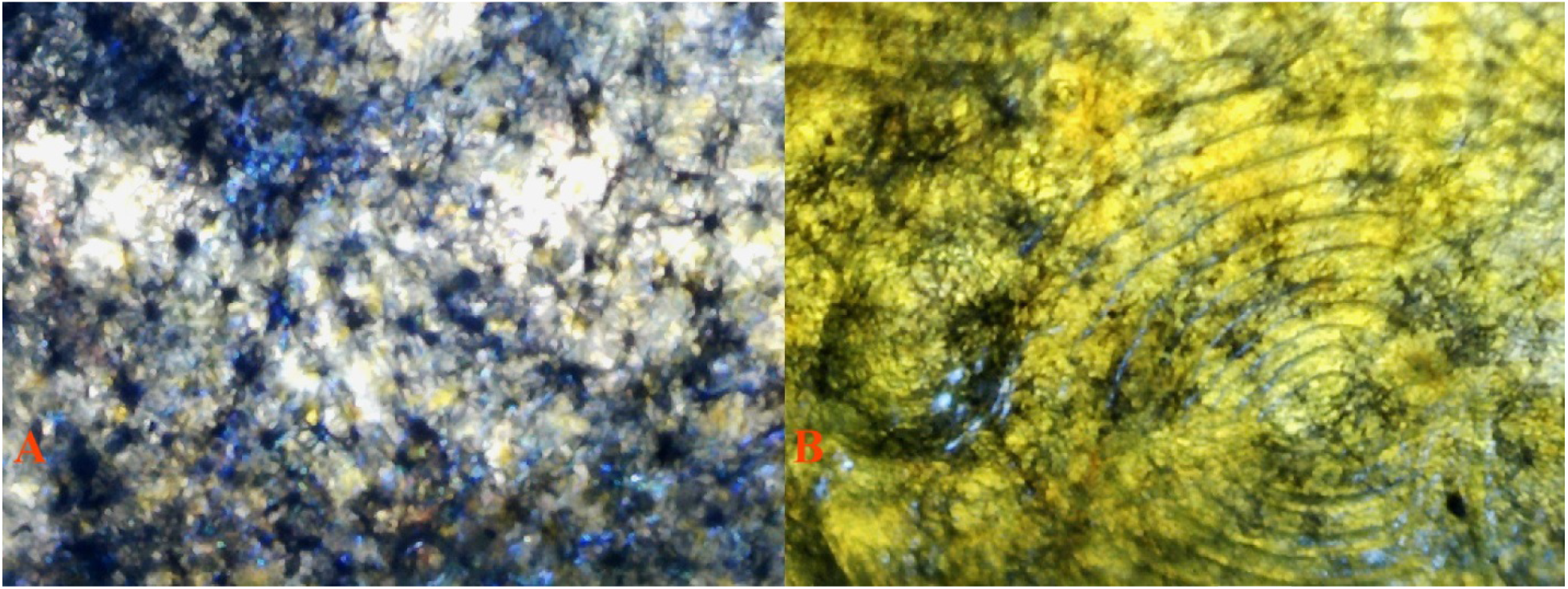
**(A)** 5 Pb 40X 16 *Pb/Pb Ab/ab (dissected)* reflected/transmitted light. Underlying violet-blue iridophore structure is clearly revealed in absence of collected xantho-erythrophores. **(B)** 4 non-Pb 40X 8 *pb/pb Ab/ab (dissected)* reflected/transmitted light. Collected xanthophores masking violet-blue iridophores in absence of erythrophores.

**Fig 14.**
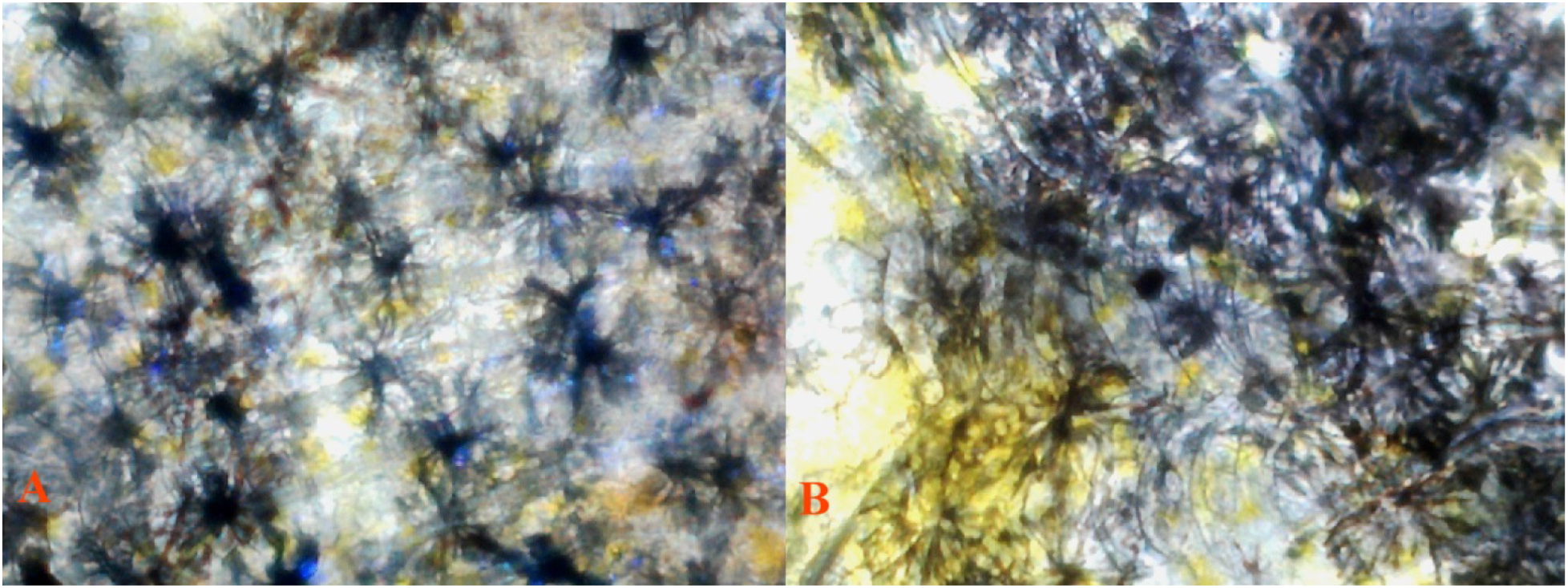
**(A)** 5 Pb 100X 1 *Pb/Pb Ab/ab (dissected)* reflected/transmitted light. Dendritic melanophore modification by homozygous Pb. Composition of dendritic melanophore-iridophore chromatophore units visible. **(B)** 4 non-Pb 100X 9 *pb/pb Ab/ab (dissected)* reflected/transmitted light. Collected xanthophores masking violet-blue iridophores in absence of erythrophores.

Collected yellow color pigment and clustered Metal Gold (*Mg* - *undescribed, see Bias 2015*) xanthophores are little affected by this erythrophore defect **(Fig 12B),** as shown in the *pb/pb Ab/ab* male. The following macroscopic photo clearly reveals near complete removal of densely packed collected yellow cells in the *Pb/Pb Ab/ab* **(Fig 12A)** male, leaving an underlying “circular ring” of violet-blue iridophores intact. As previously noted, Pb in itself has little or no effect on erythrophore populations. Albeit, Pb modification results in increased expression of violet iridophores (**Fig 12A vs. 12B**).

The macroscopic presence of underlying iridophores, lacking a xantho-erythrophore (yellow-orange) overlay in *Pb/Pb Ab/ab* **(Fig 12A)** and lacking erythrophore (orange) overly in non-Pb *pb/pb Ab/ab* **(Fig 12B)**, allows for the visual distinction between xantho-erythrophore populations. Though structural differences between xanthophores and erythrophores may be limited to variability in placement of underlying reflective crystalline platelets (Kottler 2014).

**Note:** For complete cellular microscopy study, see Bias and Squire 2017b, *forthcoming*.

## Materials

Strain ID, Breeding Strain, Description and Source (**See:** Supplemental S3 for full details).

A. Roebuck IFGA Purple Delta.

B. Shubel IFGA Green Delta.

C. Bias Ginga Sulphureus.

D. Bias Panda Moscow.

E. Bias Vienna Lower Swordtail (*Ls*).

F. Magoschitz Vienna Emerald Green Double sword (*Ds*).

G. Bias Red Double sword (*Ds*).

H. Mosseau IFGA Purple Delta.

I. Mosseau IFGA Green Delta.

J. Anderson IFGA Green Delta.

K. Anderson IFGA Purple Delta.

L. Feral Pingtung (*P. reticulata* Pingtung, Taiwan BG-2016).

M. Feral Warm Spring (*P. reticulata* Kelly Warm Springs, ID TG-2016).

N. Feral Jemez (*P. reticulata* McCauley Springs, NM TG-2016)

## Methods

Broods were raised in 5.75, 8.75 and 10-gallon all-glass aquaria dependent upon age. They received 16 hours of light and 8 hours of darkness per day. Fish were fed a blend of commercially available vegetable and algae based flake foods and Ziegler Finfish Starter (50/50 mix ratio) twice daily, and newly hatched live Artemia nauplii twice daily. A high volume feeding schedule was maintained in an attempt to produce two positive results: 1. Reduce the time to onset of initial sexual maturity and coloration, and thus reduce time between breeding’s. 2. Increase mature size for ease of phenotypic evaluation and related microscopic study.

Temperatures ranged from 78°F to 82°F. Virgin females were used in all crosses. Virgin females were obtained by removing all females from the brood tanks by “gravid spot selection” prior to male anal fins changing into gonopodia. Crosses involved breeding groups of single males (unknown or assumed genotype) or 2-3 males (known genotype) and 1-6 females. Pregnant females were isolated in breeding nets for collection of fry, and returned to breeding groups once they had produced young. All broods from each female were transferred to individual tanks and reared independently. Thus the genotypes of parental sires and dams could be verified with analyses.

## Results

A series of well-defined and structured breeding tests were undertaken during a period starting in March 2014, and ending in July 2016 upon phenotypic evaluation of final results. Efforts were made to encompass both feral populations and Domestic Guppy strains in this study.

The use of both wild-type and Domestic strains serves a twofold purpose. First, various zygosity-dependent expressions of Pb can be identified in males of either type. Second, this allows for the evaluation of natural expression of Pb modified color pigment and iridophores in Domestic females. The latter not being possible in color / tail neutral wild-type females.

Wild-type female guppies offer little in the way of visible phenotypic expression of structural color pigments. While female expression of iridophores is easily noted, this too can vary greatly between populations and strains.

The evaluation of color and tail neutral females in wild-type is limited to evaluation of increased violet-blue iridophores along the anterior lateral line. While reliable to a high degree in homozygous Pb specimens, this has not proven a reliable tool for evaluation in heterozygous Pb females.

In **Fig 15**, a pure bred male with a non-purple body was crossed to a pure bred female which was all-purple. The F_1_ were 100% Purple Body, but they were not all-purple. It should be mentioned here that all F_1_ and later generation males had some green coloration whether or not they had Purple Bodies. Therefore Purple Body is due to a dominant gene, and the all-purple phenotype involves an additional genetic component that is not dominant. It does not eliminate green coloration. Since green color is produced by light rays reflected from blue iridophores and then passing through yellow xanthophores, this shows that some xanthophores do remain.

**Fig 15.**
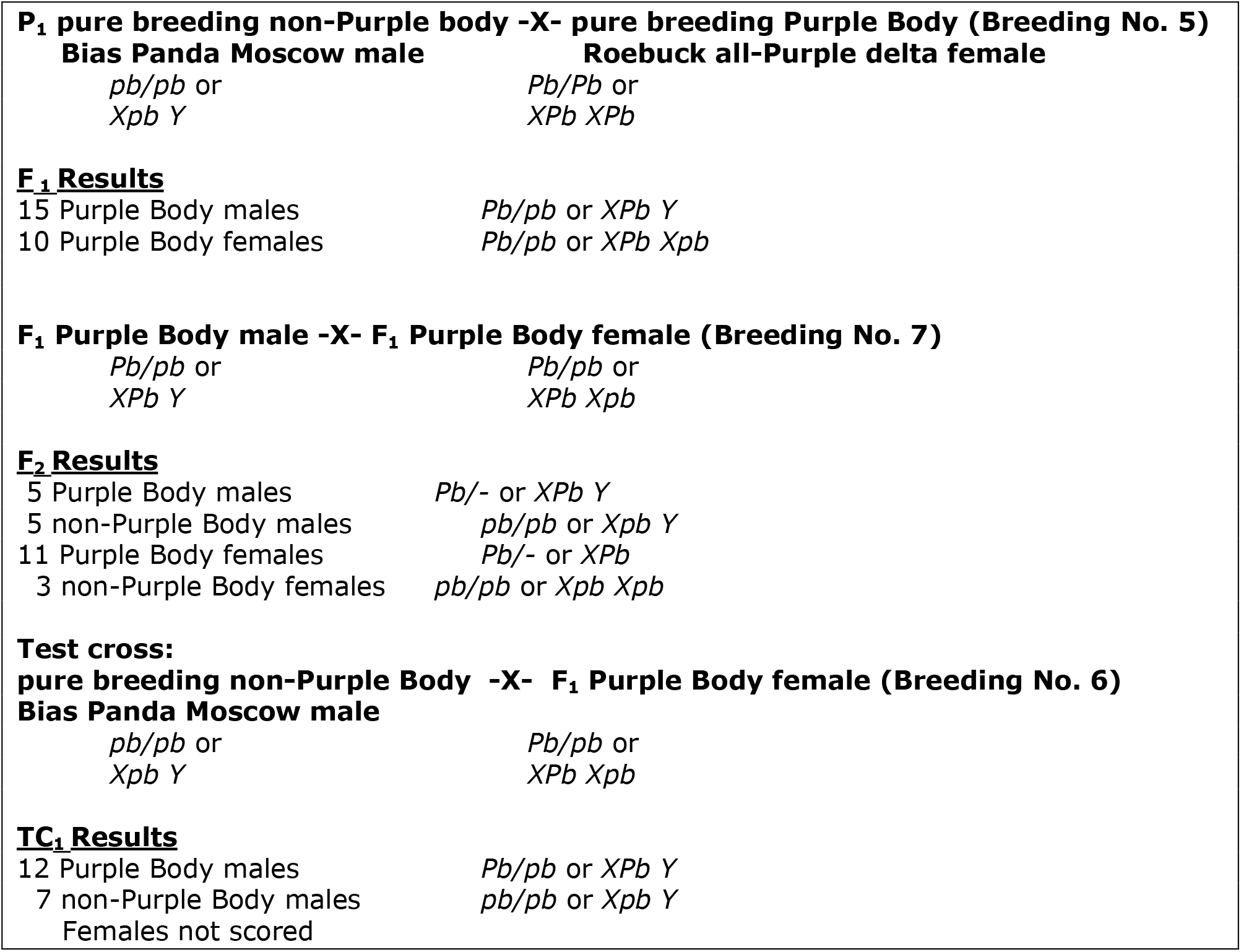
Since the P1 female had a Purple Body, Purple Body was not located on the Y-chromosome. The F_1_ and TC_1_ show that Purple Body/non-purple body is dominant and not Y-linked. The F_1_, F_2_ and TC_1_ show that Purple Body could be either X-linked or autosomal. Possible genotypes are shown.

We can eliminate incomplete dominance as an explanation with all-purple being due to a homozygous genotype and Purple Body the heterozygote with non-purple body the homozygous recessive, because homozygous Purple Body fish in the Bias Vienna Lower Swordtail strain do not have all-purple bodies. They have green as well as purple. Since the F_1_ males were Purple Body rather than non-purple body, Purple Body/non-purple body is not Y-linked. However these results did not distinguish between the possibilities that Purple Body is X-linked or autosomal.

In **Fig 16**, different F_1_ Purple Body males were crossed to pure bred non-purple body females in a test cross. The combined F_1_ ratios from several litters were 37 Purple Body: 38 non-purple body. This near perfect 1:1 ratio is expected with an autosomal dominant gene for Purple Body. In order for Purple Body to be sex-linked, the recombination frequency would have to be 50%. Although this is theoretically possible for genes located in the pseudo-autosomal region of a sex determining chromosome, it may not be likely here.

**Fig 16.**
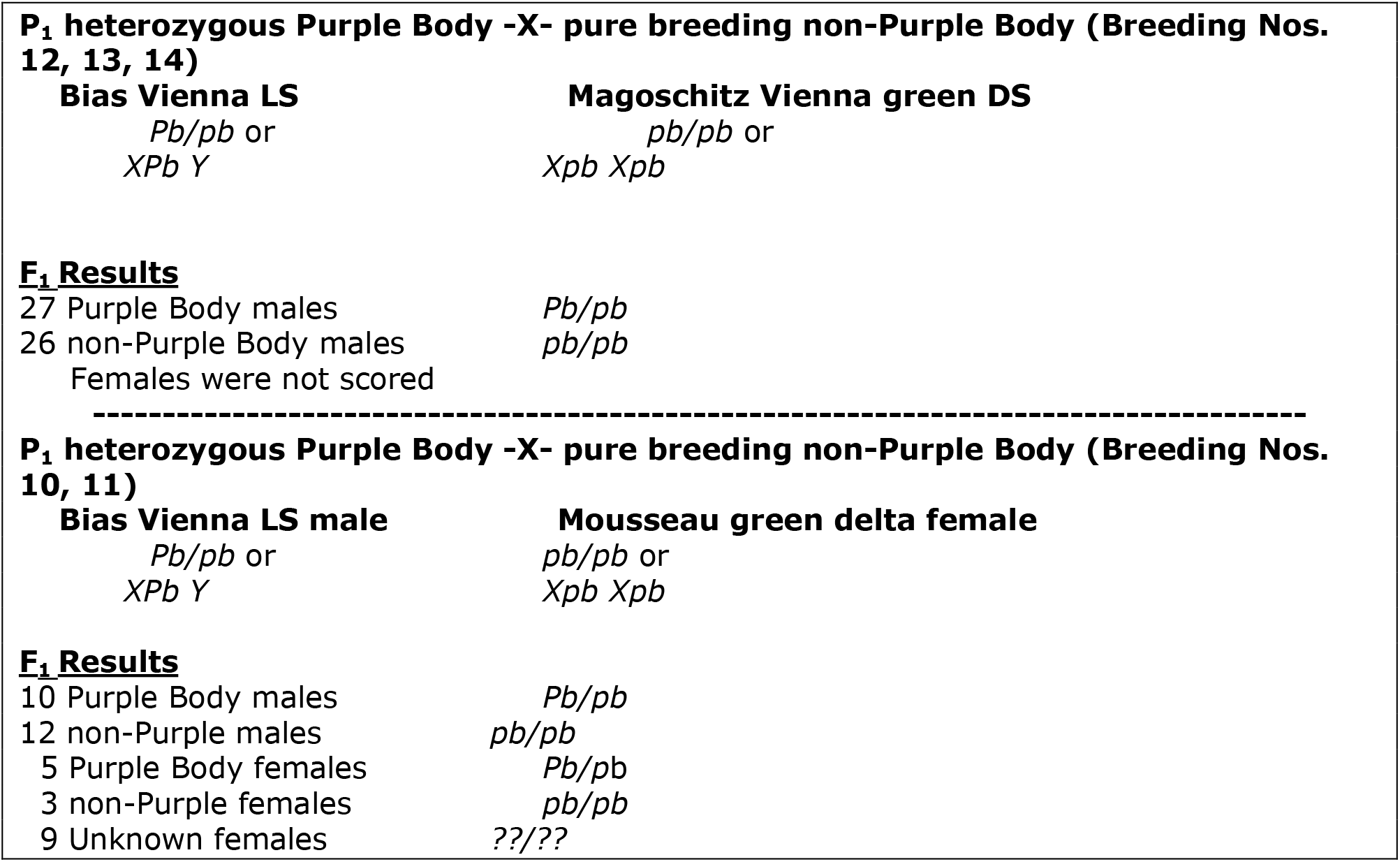
These two crosses eliminate the possibility that Purple Body is X-linked. If it were X-linked, then all F_1_ males should be non-purple body. The combined ratio of 37 Purple Body: 38 non-purple body is a near perfect 1:1 ratio, as expected if Purple Body is due to an autosomal dominant gene. Additionally, if Pb were X-linked, then all F_1_ females should be Purple Body, but instead we obtained 5 Purple Body females and 3 non-purple body females in the second cross. Therefore Pb is located on an autosome.

According to Tripathi et al. (2009), X-Y recombination is repressed (but not completely eliminated) in the pseudoautosomal region of guppies. For an additional discussion of the implications of sex chromosome structure in the guppy, see Nanda et al (2014). However, Lisachov et al. (2015) concluded that the low frequency of recombination observed by Tripathi et al. may have been due to a lack of informative markers. Lisachov et al reported a chiasma frequency of 83% of in the distal region of the sex chromosomes and suggest a “free combining region” at that end.

They compare their model of sex chromosome structure containing two “free recombining regions” on the sex chromosomes with the model of Tripathi et al and others that had one such region. All authors agree that there is a low recombination frequency in the pseudoautosomal region previously described, but Lisachov et al basically propose two pseudoautosomal regions which they call “free recombining regions”.

If Purple Body is located on the sex chromosomes, then in **Fig 16** we see 37 out of 75 recombinants and a recombination frequency of 49.3%. A chiasma frequency of 83% suggests a maximum recombination frequency of up to 41.5%. While a sample size of “only” 75 males is too small to reject the location of Purple Body on the sex chromosomes, an autosomal location seems more likely. We realize that a very large sample size would be needed to completely eliminate (or affirm) the sex chromosomal location of Purple Body. But the autosomal location of Purple Body is the most likely conclusion.

**Note:** For full description of all test breeding’s, parental and offspring photos see supplementary appendices: **S1 TABLE** – Condensed Breedings and Results and **S2 TABLE –** Expanded Breedings and Results, with photographs of examples of all fish.

## Discussion

### Purple Gene Overlooked in Previous Studies

Further study in the field should show that Pb is an integral part of wild *P. reticulata* populations, not the result of periodic population admixture through introductions. In many localized populations, especially feral ones, Purple Body phenotypic frequency may be more prevalent than non-Purple Body. It has been proposed that when a species encounters novel environmental conditions, they may adapt differently from native locales (Grether 2005). Though, general observations of *P. reticulata* wingei (*Cumana´ Guppy*) collection and study (*Fig. 3a*, Alexander and Breden 2004) suggest exclusivity of non-Pb in core area populations and predominance of Pb in overlapping areas of variant hybridization (*Fig. 3b*, Alexander and Breden 2004). This was likely biased by sexual selection preferences and compounded by environmental conditions. Thus, non-Pb is the suggested “norm” in *P. r. wingei*, while a Pb polymorphism is the “norm” in *P. r. reticulata*, with many Pb/pb heterozygotes.

Variation in predator and/or prey spectral sensitivities was early suggested as indicative of ornamentation appearing different under distinct time and space constraints (Endler 1991). While male Pb may be beneficial under specific lighting conditions as a result of female sexual selective preference, obvious questions arise. Is crypsis maintained in the presence of predators? Is Pb modification in itself an antipredator adaptation? A high incidence of Purple Body modification is noted from previously presented photographic exhibits of study specimens: (*Plate II Fig A and Plate IV*, Haskins 1970; *Figure 7*, Endler 1978; *Figure 5a*, Schröder 1983; *Figure 1*, Brooks 2002; *Figure 1* Olendorf 2006; *Figure 3*, Kemp 2008 and 2009; *Figure 2(b)* Pb Aripo River male, Millar 2012; *Figure 2*, Jourdan 2014; Pb modified “Marianne” males M14 (HP) and M16 (LP) on left, *Figure 3*, Gotanda 2014, Pb modified males “Guppy colour diversity within and between sites – five males from a given site”, Hendry Labs online 2015). Yet, the presence of the Purple Gene, and resultant implications on reduction of overall orange areas through the reduction and/or removal of xanthophores, is most often documented under “reduced expression of orange area” or “ambient light variation in hue reflection”.

## Opsin and UV

Spectral color is produced by single wavelengths of ambient sunlight. The Visible Wave Length (Perceived Color) includes: red (*620-670 nm* Bright Red / *670-750 nm* Dark Red), orange and yellow (*570-620 nm*), green (*500-570 nm*), blue (*430-500 nm*), indigo (often omitted in modern times) and violet (*400-430 nm*). Red light, with the longest wavelength and the least amount of energy, allows natural light penetration at less depth. Blue / violet light (*near-UV*), has the shortest wavelength and the most amount of energy, and allows natural light penetration at deeper depth. Violet is a true wavelength color. While Purple is a composite effect produced by combining blue and red wavelength colors.

Any visible light that penetrates the surface of a body of water is refracted; light travels slower in water vs. air. At the surface approximately one-third of surface light is scattered and absorbed by water under optimum conditions. At 1 meter up to two-thirds of the surface light spectrum is absorbed. Beyond this depth absorption has little impact on Guppy color study as it is beyond the zone of habitation. Environmental conditions of natural Guppy habitat are varied and under diverse lighting conditions. At mid-day nearly all light may be absorbed by water, and little is reflected. While in the early morning and late afternoon a reduction of absorbed light is seen, and more is reflected. Absorption of both sunlight, and its energy, may be further reduced by cloud cover, plant cover, time of day, time of year, altitude, water molecule structure, water flow, and turbidity.

With such a broad range of spectral irradiance, it is only natural that study of Guppies would evolve with consideration of visual capabilities extending from near-UV to far red. Vision is based on retinal (forms of Vitamin-A) bound to opsin proteins. Visual receptor cells in Guppies contain light sensitive opsins along outer segments. Opsin “pigment pairs” contain two pigments with different wavelength peak sensitivity; one rhodopsin (also known as visual purple) and one porphyropsion (Bowmaker 1990). *Poecilia reticulata* possess at least six (see later study indicating nine) long-wavelength sensitive (*LWS*) opsins, as compared to other teleost species averaging two. Study has shown that *reticulata* opsin divergence occurred both pre and post divergence from a close related species, Lucania goodei (Weadick 2007). Results favoring four out of six opsin duplication events occurring after divergence, likely playing a role in *P. reticulata* extreme polymorphic diversity in comparison to L. goodei.

Variation in colors and color characteristics such as hue, depth, etc. cannot be important in female based sexual selection unless the female can detect these color characteristics. Therefore the evolution of color characteristics must be accompanied by the evolution of the ability to detect the colors. Endler showed that selection for spectral sensitivity variation in both short-wavelength sensitivity (*SWS*) and long wave sensitivity (*LW*S) is due to an hereditable factor (Endler 2001). Recent study indicates color vision varies across populations, and that populations with stronger preferences for orange had higher LWS opsin levels (Sandkam 2015a and 2015b). It has been shown that Guppies are able to perceive UV wavelengths, and that males reflect UV from both structural color and color pigment with variability between individuals. It was further shown that female association preference with males is under long wavelength (*UV-A*) conditions in which orange is visible (White 2003). While this study was suggestive of females having little or no sexual selection preference for either low UV or high UV males, it did not specifically focus on benefit derived from reflective qualities of Pb under reduced ambient lighting conditions.

Further Opsin studies of *Poecilia wingei*; i.e. the Cumana’ Guppy, reveal that gene duplications have increased the number of opsin genes and one additional opsin gene (*LWS A180*) is the result of divergence from an ancestral poecillid gene, rather than duplication. The different rod and cone classes produce a wide sensitivity to different wavelengths of light in the Cumana’ Guppy (*Poecilia reticulata wingei*). Variations in synthesis of these opsins may generate the diversity of types of color perception found in variant populations of Guppies (Watson 2010). Male polymorphism in Guppies is, in part, the result of an expanded number of opsin genes, which help foster female sexual selective preference for orange color spotting and often diversity (rarity) of pattern. Interestingly, a molecular level study (Ward 2008) showed the bulk of LWS mRNA in Cumana’ was derived from the LWS A180 gene. No mention was given of the presence of SWS in Cumana’, a population in which Pb is typically non-present in core area populations. Though, presence of underlying violet iridophores is expected. The expansion of Watson’s studies to other Guppy populations with different color characteristics seems warranted.

Teleost species possessing fluorescent color pigment have the capability to absorb reduced available light, and re-emit at long wavelength. This process, known as bioluminescence, is produced through a chemical process and is often restricted to deep water marine species. Study of those emitting red luminescence has shown that mechanisms utilized in long wave fluorescence involves the collection and modulation of overlapping dendritic melanophores and motile dendritic red iridophores in “chromatophore complexes” (Wucherer 2012, 2014). Individuals expressing Pb exhibit higher violet to blue iridophore density. Whether this results from increased cell population levels or simply the result of increased visibility from reduction of yellow xanthophores has not been determined. Existing red erythrophore populations appear unaltered.

While bioluminescence is not known to be applicable to *P. reticulata*, microscopic results (Bias, *unpublished*) in both Pb and non-Pb are suggestive of the aggregation of motile melanophores in conjunction with the collection of blue and violet iridophores among, between and beneath dendritic structures, forming similar chromatophore units **(Fig 17)**. A similar association has been shown in general imagery of recent research (*Fig. 1 and Fig. 6c*, Kottler 2014). Whether this represents increased population numbers or density, or simply increased visibility in conjunction with guanine crystalline platelets was not determined.

**Fig 17.**
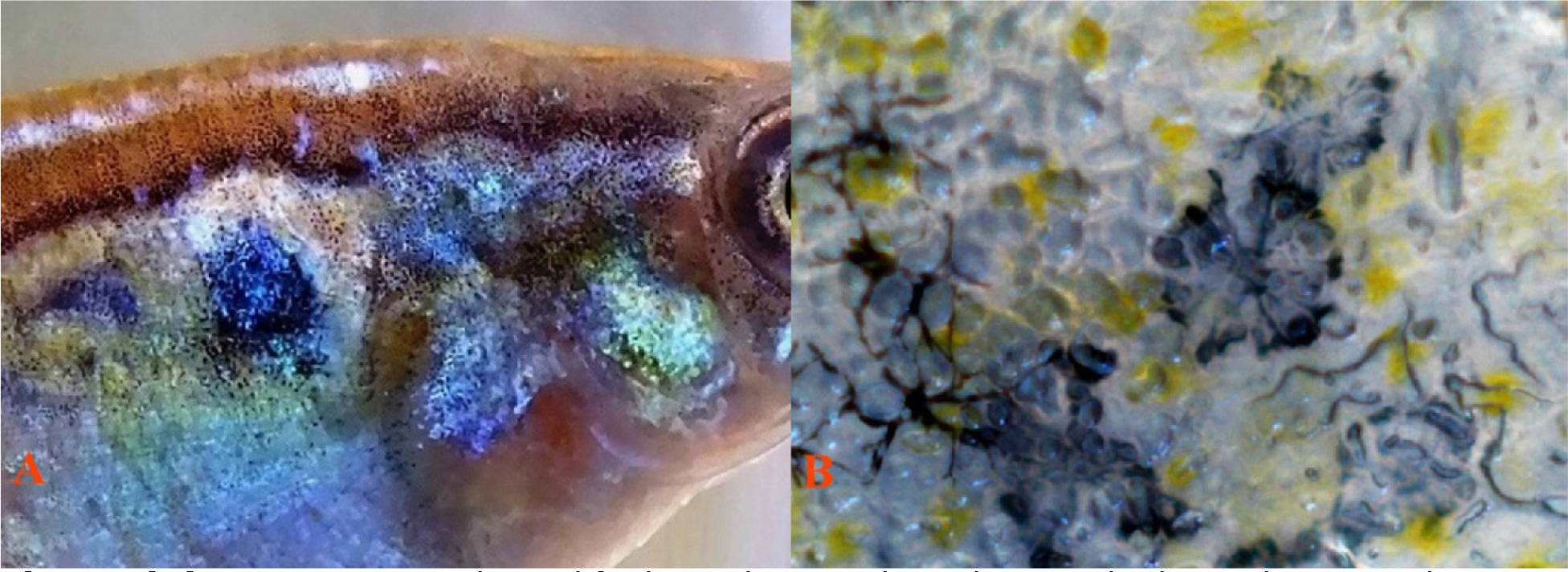
**(A)** Heterozgyous Pb modified Dendritic melanophore-iridophore chromatophore units. (photo), and **(B)** Non-Pb Dendritic melanophore-iridophore chromatophore units. (100X non-dissect, reflected lighting).

Goda (1995) reported the existence of what appear to be minimally-reflective blue chromatophores (cyanophores) in two marine species; to this date no such dendritic blue color pigment cells were identified in *Poecilia reticulata*. Nakajima (1999) in a brief report noted “dendritic “bluish-white” chromatophores” with reflective qualities, observable only by means of a fluorescence microscopy. He stated these structures could not be seen with standard reflective or incident light microscopes. This was found only in one strain (an undescribed type of *albino*) out of 17 strains studied; he suggested the result of genetic polymorphism, i.e. strain specific and not species specific.

## UV Vision and Mate Choice

An inaugural study into UV vision and mate selection concluded female preference for UV+ males, and males slightly preferring UV− females (Smith 2002). In conclusion, it was thought hue discrimination vs. brightness was the primary positive benefit of UV capabilities. While this study was based on sexual selection preference in matings, non-sexual preference origins based on food color detection have been shown (Rodd 2002). Prior studies heavily focused on the shallow water habitat of Guppies, subject to UV radiance from the sun. None had attempted to assert a direct correlation between UV reflected color, pattern and vision, mate choice, and individual preferences. Several of these earlier studies re-affirmed courtship activity was at its highest during dawn and dusk. These are periods during which SWS and LWS are visible from low angle ambient sunlight (Endler 1987, 1991, 1992; Loew 1990).

Smith concluded that there was no discernable UV preference between early morning simulations 1-3 hours after lights on or 1-6 hours, with 1-3 being indicated as traditional testing time by researchers (Smith 2002). This is worthy of note for at least two reasons: 1. As previously noted, yellow color pigment in Guppies is highly motile (subject to constriction of yellow pigment cells and ectopic melanophores). Expression is often based on hierarchal ranking and further influenced by courtship display. Therefore, during early morning hours actual expression of yellow is commonly reduced or near non-existent after long periods of darkness. This alters general expression of brightness in xantho-erythrophore orange spotting and yellow ornaments in early hours. In turn, this constriction temporarily increases the visibility of underlying reflective structural color and pattern (Bias, *unpublished observations and data*). 2. If present, Pb as one of its primary modifications further removes yellow color pigment cells and increases visibility and/or population levels of UV reflective violet iridophores, further altering previously mentioned expressions in both early morning and late afternoon hours.

*P. reticulata* have been shown to possess transparent ocular media (Douglas and Hawryshyn, 1990), making them sensitive to UV spectrum in the absence of visual pigments in cones. While little is known in regard to *reticulata* transparent transmission, some species of mammals have been shown with sensitivity in the 320-340 nm range (*see citations*, Douglas 2014). Several studies confirm the presence of UV-sensitive retinal cones, UV-transmittable ocular media, and SWS opsin genes in Guppies (Douglas 1989, 1990; Archer 1987, 1990; Weadick 2007; Ward 2008; Watson 2010; Smith 2002).

While Archer was unable to prove the existence of visual pigments extending into the accepted starting range for peak sensitivity (*maximum absorbance* - λ_max_) in UV spectrum (*UVA 380-400nm*), he showed well marked clusters at λ_max_ 410nm, 465nm and 573nm. He concurred with earlier studies asserting Guppies are polymorphic for color vision in LWS, with most rhodopsin-porphyropsin polymorphism in cones absorbing yellow, orange and red. Kemp in turn reported UV reflectance in violet-blue iridophores and orange spots ranging from 350-400nm (Kemp 2008). A molecular level study (Ward 2008) indicates a higher than normal duplication and divergence of 4 distinct LWS in *Poecilia*, as compared to other species. Expressed concurrently, they provide the means for both increased SWS and LWS discrimination.

Temperature has been shown to have direct effect on corneal-positive deflection in dark-adapted spectral sensitivity under varied wavelengths in Zebrafish. The result was a shift in rod visual pigments. Cold water (*22-25° C*) spectral sensitivity resulting from a mixture of rhodopsin-porphyropsion, and warm water (*22-25° C*) rhodopsin. Under both conditions contributions were made by UV cones at λ_max_ 362 nm (Saszik 1998). Seasonal variation in composition of rhodopsin-porphyropsin levels has been shown in Masked Greenling (*Hexagrammos octogrammus*) resulting in resulting in significant λ_max_ shift to LWS (Kondrashev 2008). It is conceivable that similar fluctuations in Guppy vision may occur under seasonal lighting and temperature changes in the wild, and fluctuations under laboratory study. If so, these fluctuations may elicit distinctly visual responses, in Pb and non-Pb wavelength discrimination for breeding preferences. Similar consideration should then be given to predatory response systems.

With the discovery of variation in opsin expression of the Guppies nine opsin cones between individuals, it has been suggested that new designs in behavioral study are warranted in regard to mate choice (Rennison 2011). Modification of iris pigment is noted in Pb, resulting in increased levels of violet iridophores, near predominance, as compared to non-Pb. A similar situation is also found with the modification by other traits, such as Metal Gold (*Mg*) (Bias, *unpublished breeding notes*), producing not only proliferation of reflective yellow color pigments in the body, but also in the scleral ring and iris. Developmental changes suggest enhanced wavelength discrimination in adults as compared to juveniles (Laver 2011). Modifiers of corneal dorsal / ventral oriented pigmentation in both males and females, such as Pb and Mg, may produce results which act as “corneal filters”, not limited to the seasonal restrictions previously discussed, in fostering SWS and LWS discrimination.

## Predation

A study shows that environmental light has an effect not only on spectral sensitivity based on level of maturity, but also coloration independent of age (Hornsby 2013). Greater risk from predation occurs during periods of highest environmental lighting conditions (Endler 1987). Our supposition is that the primary benefit of erythristic Pb modification **(Fig 18A-B**) is derived from low light conditions; either in heavily forested upland canopy and/or during periods of low angle sunlight in the early morning and late afternoon. This is when reflective qualities of Pb modifications are likely to become highly visible in the UV and/or near-UV spectrum **(Fig 18A)**, even though to the naked eye both Pb and non-Pb color is nearly identical during this time. While in open canopy and/or during periods of bright high angle sunlight non-Pb appears “brighter orange”, as compared to Pb appearing “darker orange” **(Fig 18B)**, and is favored as a sign of male fecundity.

**Fig 18.**
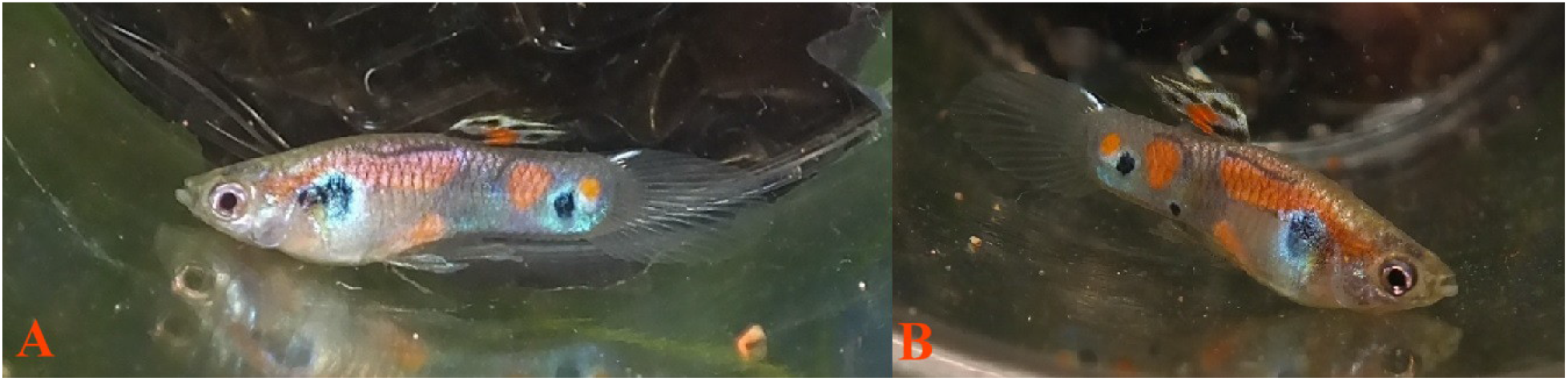
**(A)** Pb male expression with low angle ambient light source appearing modified pink, **(B)** Pb male expression, in same individual, lacking low angle ambient light source appearing dark red.

In a study of the Trinidadian Pike Cichlid (*Crenicichla frenata*), it was noted *Poecilia reticulata* males generally express not only reduced polymorphism, but also reduction in the intensity of color and pattern. Results indicate that predation imposes restriction on male ornaments expressing patterns comprised of blue and iridescent structural color spots (Endler 1980; Kemp 2009). Millar also suggests the results indicate that teleost predation favors less orange spotting with smaller size, with an indirect effect by local predatory shrimp (Millar 2006). In follow-up Millar generally finds, *“high-predation males typically have more relative UV reflectance when all rivers and sites are considered in the same analysis”* (Millar 2012), further showing UV as a private signal in some populations, but not all. Worthy of note, this and several other studies postulate that other factors are contributing to selection beyond a simple high-low predation contrast (Millar 2006; Karim 2007). Pb modification of both ornaments and overall coloration should be considered in such future studies.

Weadick asserts in his discussion, *“establishing that one of the guppy’s most dangerous predators can detect a wide swath of the spectrum represents a critical step toward fully understanding the nature of selection on color patterns in this system* [*C. frenata* -*P. reticulata* predation]” (Weadick 2012). Citing earlier studies, the author cautions against interpreting results, based on limited LWS / SWS receptor sensitivity studies, involving effects of predation on prey color and pattern. Limited or single samples do not rule out variability in visual receptors in a single species of predator in multiple locations within its range.

Of five opsins isolated in more recent *reticulata* studies, two were identified as “maximally sensitive” in producing SWS vision. Of more importance in consideration of potential Pb biotic benefit from modification, it was found that *C. frenata* was notably “insensitive” to UV light, being unable to discriminate hues in the lower part of the visual spectrum (Weadick 2012).

The predatory prawn (*Macrobrachium crenulatum*) has been shown capable of UV discrimination; prey individuals expressed higher degree of melanophore spotting and reduction in levels of reflective iridescence (Endler 1978; Rodd 1991; Kemp 2008). Several studies indicated no reduction in the amount or size in orange spotting, as piscine predators are generally unable to detect Long Waves (Endler 1978; Houde and Endler 1990; Kemp 2008). Generally, this indicates the absence or reduction of Pb modification for SWS UV visibility in the presence of *M. crenulatum*. While the color photo quality (*Fig. 7g*, Endler 1978) of Paria population makes it difficult to verify presence of Pb, it is suggested. Conversely, the photo quality (*Fig. 3*, Kemp 2008) shows large orange spot modification from Pb in the Marianne population and predominance of Pb modification in the Quare population. The author noted, “relatively low UV peaks in Marianne fish colours are notable given that this population is the only one to co-exist with a strongly UV-sensitive predator *(M. crenulatum)*”.

To avoid predation, *M. crenulatum*, is primarily a nocturnal feeding species (Bauer 2011) that in turn preys heavily on *P. reticulata* (Magurran 2005). Orange spots on the pinchers were shown to act as a diurnal sensory lure in attracting Guppies to the anterior head region of the prawn. This sensory bias was successful to a higher degree in allopatric *reticulata* populations, and to a lesser degree in sympatric *reticulata* populations (Rodd 1991; De Serrano 2012). While increased predation on any age class of individuals will have an impact on the overall constitution of color and pattern, the overall effect of predation on Pb modification of said pattern will be reduced by an autosomal dominant mode of inheritance that is transmitted by either sex. This, being in direct contrast to early opinion that “anything except predation has been important in the evolution of the orange colouration” (Rodd 1991).

Hughes, et al., (2005) raises the question of to what degree does balancing selection contribute to the maintenance of color pattern and size variation found within a population or across populations? Statistical analysis of sib x sib breeding results structured to identify Genetic-by-Environment interactions (*G x E*) effects on color and size were undertaken. Results showed that while dietary intake did have the expected influence on size, it did not have any detectable influence on color and pattern. But they did not consider Purple Body coloration in their study.

According to Hughes, three mechanisms had been previously put forth at the time to explain the maintenance of these polymorphisms: 1. Gene flow between populations with differing selection regimes (Endler 1980), 2. Negative genetic correlation between male survival and attractiveness (*antagonistic pleiotropy*) (Brooks 2000), 3. Negative frequency-dependent selection (Farr 1977, 1980b; Hughes 1999). These proposed mechanisms are not mutually exclusive.

This leaves us with a bit of a dilemma, as these previous studies dealt with variations in color and/or pattern type and size. The resultant benefits of Pb on mating success likely have little specifically to do with male rarity or novelty (Farr 1977) since Pb is often present at high frequencies in populations. Rather they derive primarily from increased near-UV visibility under specific ambient lighting conditions (time of day) in conjunction with water conditions (turbidity and/or chemistry) and locale (upper and/or lower drainage, canopy cover or lack thereof).

The presence of X-linked and/or autosomal coloration in females (limited to *xantho-erythrophores* and *melanophores*) has been shown through testosterone treatment in high-low predation sites (Gordon 2012). High predation females consistently showed less coloration compared to low predation. Expression of homozygous autosomal dominant Pb, similar to identified autosomal recessives found in *Poecilia reticulata*, is a modifier of total existing body color and pattern pigmentation (*xantho-erythrophores, structural colors and melanophores*) in both males and females.

It is an autosomal trait capable of modifying extent color and pattern found in a population and across populations, acting as “a permanently protected reservoir available in the female population in which, whether they are present in heterozygous or homozygous condition, they are sex limited and will not be phenotypically expressed” (Haskins 1951). The Pb gene is partly protected from strong selection by its presence in females, where it is minimally expressed. The Haskins’ study set out to, “ascertain, in a very general way, whether X-linked and autosomal linked color patterns occur in wild populations”. While not identifying any specific traits, they concluded in their discussion, “data… …indicates that this in-deed is true”.

Here we have the Purple Body (*Pb*) gene identified as the first autosomal gene to be described as existent in high frequencies in both wild and Domestic Guppy populations; one capable of pleiotropic effects on all existing color and pattern elements at multiple loci. Several questions arise. Firstly, “Should Pb with an autosomal mode of inheritance, in itself, be considered a mechanism capable of balancing overall color and pattern polymorphisms?” Secondly, “While positive in overall benefit under specific conditions, does the Purple Body condition present a detrimental pleiotropy to the individual, especially in homozygous condition?” These questions seem worthy of further study in controlled lab settings and under natural conditions.

Studies continue to focus on fitness through heterozygosity maintained and limited to male ornamentation through multilocus heterozygosity (*MLH*, Herdegen 2014). Studies on female ornaments in “wild-type” are few if any. This is likely based on the assumption that wild-type *P. reticulata* females are essentially “color-trait neutral” as a form of camouflage, with expression of pigmentation limited to Domestic Guppy strains (Goodrich 1944; Lindholm 2002; Kottler 2013 and 2014), expressing limited coloration to the naked eye. Yet, current microscopy reveals female coloration is not solely limited to counter-gradient expression for the benefit of camouflage.

Females possess all classes of chromatophores, and are subject to Pb modification for expression of near-UV reflective qualities as well. MLH was correlated as a “significant predictor” in relation to overall area of spotting; it was not linked to total numbers of spots. The end results favored “genome-wide heterozygosity” based on individual markers for spotting identified as non-heterozygous. The potential immunologic benefits of heterozygosity have long been subject of study; the major histocompatibility complex (*MCH*, Yamazaki 1976; Brown 1997), the parasite-resistance hypothesis (Hamilton and Zuk 1982). A direct correlation has linked to heterozygosity and male fecundity in the form of motivation and increased sperm counts (Mariette 2006; Zajitschek and Brooks 2010). The same correlation is known to result in a higher degree of female fecundity. Heterozygous Pb is once again worthy of prime consideration.

In a study of Betta splendens (Clotfelter 2007), evidence indicated that purple males derive a greater immune response than do red males. When both color morphs were supplemented with additional carotenoids, purple males diverted fewer carotenoids to color maintenance and exhibited greater immune response. “Unlike other species in which this trade-off has been examined, where the ability to maintain carotenoid-based coloration is condition dependent and results in a range of red and less-red phenotypes, male B. splendens have genetically determined color morphs… …Redder individuals (positive PC2 [sic *600-70 nm*] values) provided with supplemental carotenoids showed an increased inflammatory response to PHA [sic phytohemagglutinin] and greater redness, whereas bluer individuals (negative PC2 [sic *320-520 nm*] values) showed no change in coloration and instead mounted an even greater immune response.” They suggest that red males require higher carotenoid concentrations and are genetically oriented to load balance stockpiled reserves between color maintenance and health issues.

Is heterozygous Pb selectively favored in males while homozygous Pb is unfavorable? If so, then balancing selection would tend to maintain both alleles in the population. If the heterozygote is greatly superior to the two homozygotes, then selection would tend to maintain both alleles balanced in the population. If for example the initial allele frequencies were 0.5 Pb and 0.5 pb that would generate 25% Pb/Pb, 50% Pb/pb, and 25% pb/pb in the next generation in the absence of selection. But if selection eliminates most of the Pb/Pb males and perhaps some of the pb/pb males as well, then the Pb/pb males would be selectively favored in each generation due to heterozygote superiority. The final frequencies of each of the two alleles and three male genotypes would depend upon the sum of the selective pressures on each of the individual genotypes, and these might vary significantly over time and locality.

## Feral Populations

Studies of feral Guppy introductions are often considered of lesser value in comparison to those done in native locales. While of value from an evolutionary standpoint in subsequent studies, they often lack historical founding population references. Yet, in many ways these events are no different from natural colonization by single individuals or limited numbers. Intentional feral introductions, of known parentage, have been performed in several locations and are thought to offer a more complete comparative analysis resulting from environmental change. As previously stated such introductions, seemingly unknown to researchers, are often revealed by analysis to share a common characteristic: the presence of Pb **(Fig 19)**.

**Fig 19.**
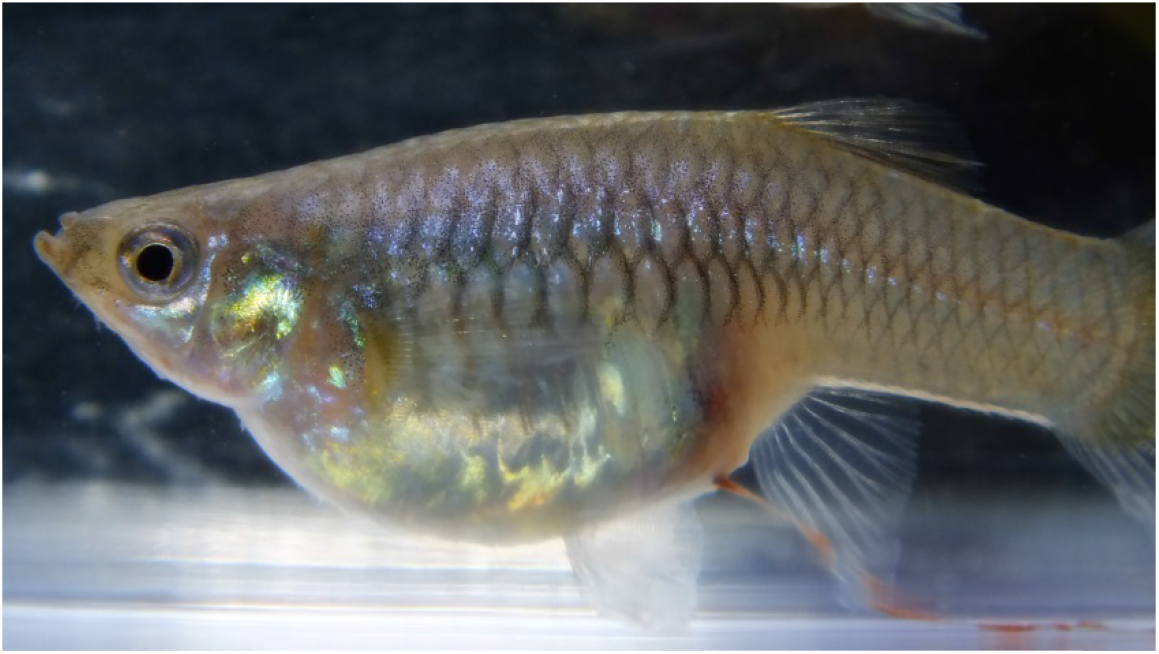
Homozygous (*Pb/Pb*) wild-type female.

One intentional feral introduction, considered of value by weight of published studies, involves the stocking of high predation Yarra River (Trinidad) stocks into low predation Damier River (Trinidad), with later colonization of high predation Damier. Attempts to capture expression of orange and reflective qualities of populations under variant ambient lighting have been made (Karim 2007; Kemp 2009). Observations were limited to orange, black and blue and total amount of pooled color. The Damier population (high and low) showed no divergence in orange values as compared to the foundation high predation Yarra population. A rapid increase in survival fitness traits has been shown to be comparable to that of captive reared wild stocks (Gordon 2009). Final analysis in both studies was based on commonly used statistical measures.

Here again, the presence or absence of Pb within the overall population was not directly studied. Pb has been demonstrated to have potential implications on the divergence through modification of existent orange and structural coloration.

Domestic Breeders have demonstrated that specific genes for many expressions of female ornaments are contained in the genotype through sex-linked (possibly in the form of initial cross-over events) and autosomal modes of inheritance. Still other traits stem from new mutations or recombination in long-term captive bred stocks (domestic and wild). Many are known to persevere in both controlled breeding’s of wild and feral populations (*Turure high-low controlled feral introduction*, Gordon 2012). The study of non-expressed female coloration (wild and feral) by Gordon was limited to testosterone treatment for xantho-erythrophore color pigment and melanophores. It is erroneous to assume that wild and feral females do not express variability in violet-blue structural iridophore ornamentation in limited form, easily discernable to the trained naked eye. The Purple genotype is now demonstrated to be a potential female study marker in heterozygous and homozygous form, in both wild-type and controlled breeding’s.

Studies of wild and feral populations fail to support a correlation between in-breeding and increased level of spotting (Zajitschek and Brooks 2010). Similar color patterns have been found in wild and introduced feral populations, in both high and low predation locales, each expressing no measureable differences in either number or size of spotting (Martínez 2016). Orange spotting, in controlled laboratory settings, has been shown to reflect increased expression with a higher inbreeding co-efficient (Nicoletto 1995; Sheridan 1997; van Oosterhout 2003; Mariette 2006). A similar observation is found in the tanks of Domestic Breeders through artificial selection, as evidenced by many diverse carotenoid xantho-erythrophore (yellow-orange) and pteridine (red) phenotypes, in spotted and solid expressions. The Purple gene has now been identified as having the ability to modify extent genome-wide yellow-orange-red color and pattern spotting in heterozygous and homozygous fashion.

Female preference consistently fosters sexually selected traits across populations in selection for males exhibiting similar phenotypic traits (Kodric-Brown 1985). Xantho-erythrophores are considered indicators of male over-all fitness and therefore are attractive to females. Diet has been shown to increase brightness of existing orange spotting, but not size of spots (Kodric-Brown 1989). Orange carotenoid pigments have been shown to elicit positive response initially at a distance and catch the female’s attention. While the importance of structural violet-blue iridophores take precedence during actual courtship and retain the female attention (Endler 1983, 1984; Kodric-Brown 1989 and 1996).

Kodric-Brown (1989) utilized two study populations, Paria wild [*Trinidadian*] and Jemez feral [*McCauley Springs, NM*]. Jemez feral express reduced orange spotting in comparison to other populations. Not surprisingly, it was reported “Females from the Jemez population showed significant disagreement in their preferences for individual males.” Jemez males exhibit increased iridophore spotting (blue and white), and increased circular and linear melanophore spots. Jemez males are very iridescent violet-blue **(Fig 20-21)**, often expressing: Iridescens (*Ir*), (Winge 1922b; See also: *Iridscens (Ir)*, Blacher 1928; *Smargd Iridescens (SmIr)*, Dzwillo 1959; *Blue Iridescent Spot*, Kottler 2013; *Reflective Dorsal Spot (RDS)*, Bias 2013).

**Fig 20.**
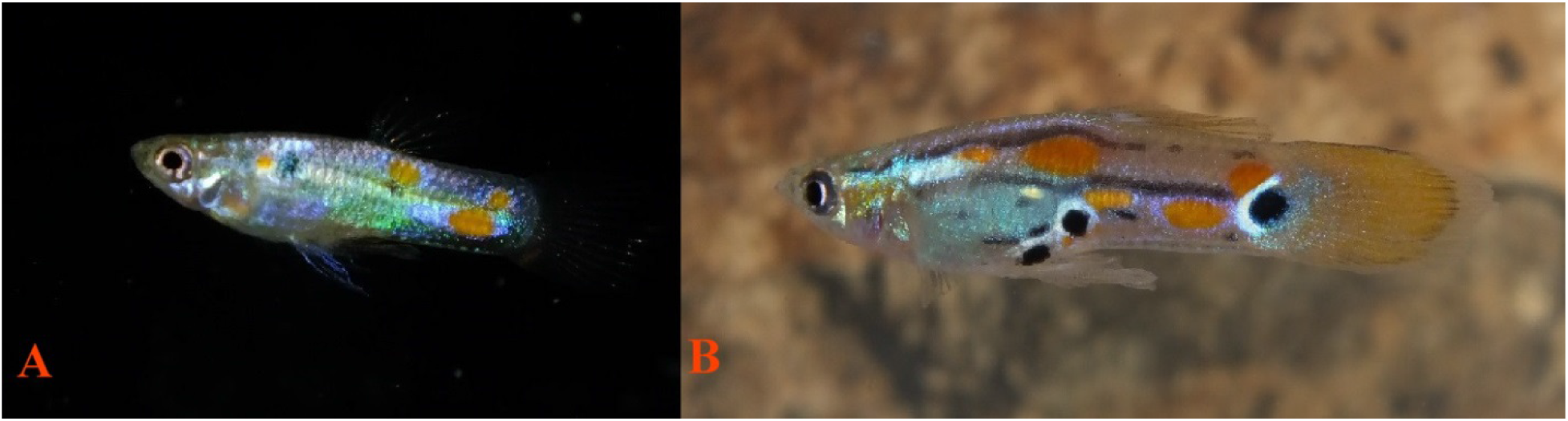
**(A-B)** Wild-caught Pb Jemez males, McCauley Springs, NM (2016 Collection), dark spotting ornament expression lacking low angle ambient light source.

**Fig 21.**
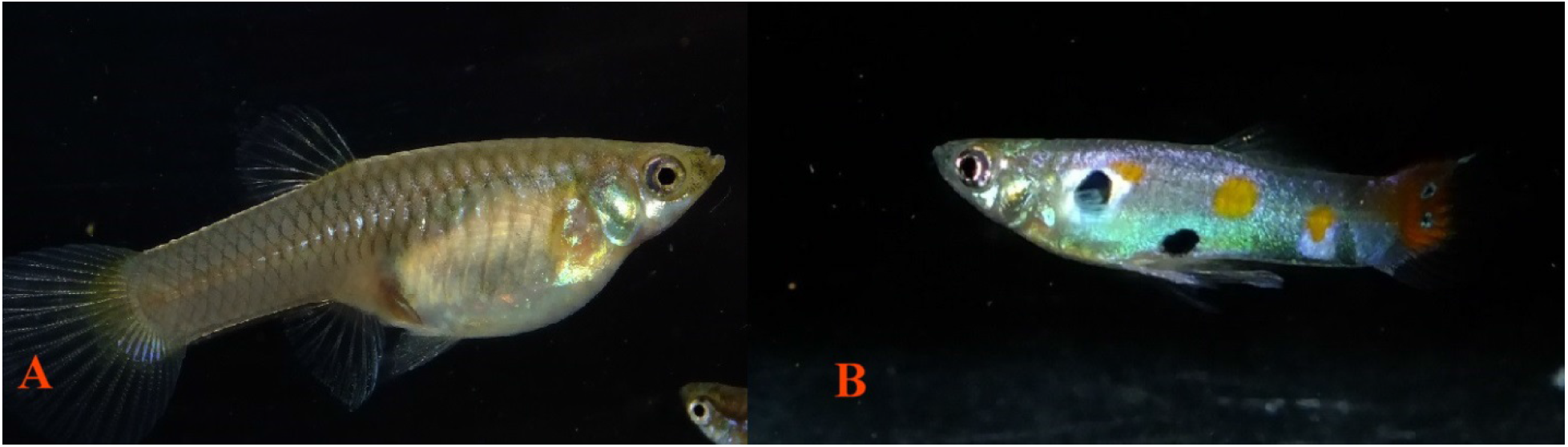
**(A)** Wild-caught Pb female expressing Pb modification in caudal base, and **(B)** wild-caught Jemez male, McCauley Springs, NM (2016 Collection), dark spotting ornament expression lacking low angle ambient light source.

The majority of Jemez individuals, males and females, recently collected and reared in subsequent captive breeding are modified Pb (Bias, *unpublished breeding notes*). It is assumed that there is sexual selection for autosomal Pb modification in both males and females because of increased UV visibility interacting with violet-blue iridophores. A recent study confirms that females do have strong preference for UV reflective color pattern in males, though male preferences were omitted in the study (Kodric-Brown 2002). Jemez males expressed the majority of their iridescent colors below 400 nm (*near-UV and UV* spectrum).

Kodric-Brown (2002) stated: “Among the iridescent colours, white and purple strongly reflected below 400 nm, but green and blue did not. Gold (yellow-orange) also showed a UV component. Generally, the area of UV reflectance closely matched the area of iridescence visible in longer wavelengths… …Although the overall colour pattern was the same when viewed in the visible and the short wavelengths, certain aspects of the pattern were more noticeable in the UV wavelengths. Melanin (black) spots surrounded, either completely or partially, by a ring of iridescent white, and gold spots next to black areas provided a striking contrast in the UV.” As previously shown melanophores spots are known to reside in close proximity and intermingled with violet-blue iridophores.

The obvious benefits of increased visibility and/or numbers of violet iridophores through Pb modification are again asserted. Guppies are known to have multiple UV sensitive receptors, as opposed to native predators that are UV insensitive with a reduced number of receptors in short wavelengths. Feral populations, lacking reduced predation, such as Jemez, take advantage and express increased incidence in all areas of reflectivity, including “white patches”. The latter being highly UV reflective, but also visible in shorter wavelength and subject to predation in wild populations (Kodric-Brown 2002). Pb as an autosomal modifier of the existing phenotype, has now been shown to increase some chromatophore populations and/or visibility in the UV spectrum. This in part, proves earlier research suggesting, “Selection should favour signalling in these short ‘private’ wavelengths” (Endler 1991).

## Summary and Conclusions

From what can be ascertained, the bases of cellular studies continue in attempts to quantify orange and structural color and/or pattern values into measureable form, without taking into account two distinct genotypes found in wild and feral populations: Pb and non-Pb. Yet, recurrent photographic evidence such as provided by Millar and Hendry [*Fig. 2 UV (a) and colour (b) of a male guppy from the low-predation sampling site in the Aripo River]* (Millar and Hendry 2012), and Gotanda and Hendry [*Fig. 3, M14 (HP) and M16 (LP)*] (Gotanda and Hendry 2014), clearly exhibit UV-enhanced modification by the Purple gene.

Crosses between Purple Body and non-purple body guppies prove that the Purple Body gene is not simply Y-linked. Although the results did not completely rule out the possibility that the Purple Body gene is located in a frequently recombining pseudo-autosomal region of the sex chromosomes, that interpretation is considered to be extremely unlikely. The final conclusion is that the Purple Body gene is located on an autosome, with an incompletely dominant mode of inheritance.

*Poecilia reticulata* exist in a heretofore undocumented polymorphic state; Purple Body (*Pb*) and non-Purple Body (*non-Pb*). The co-existence of the two phenotypes suggests a selective advantage under predation (crypsis) and in sexual selection (conspicuous pattern) under diverse ambient lighting conditions.

The violet-blue chromatophore unit and removal of xanthophores by Pb modification is required to produce an all-purple phenotype. The Purple gene has the ability to modify extent genome-wide chromatophore populations in heterozygous and homozygous condition, with increased visibility in the UV spectrum. As a result, this demonstrates selection favoring short “private” wavelength signaling.

Pb is now identified as the first polymorphic autosomal gene to be described as existent in high frequencies in wild, feral and Domestic Guppy populations. It is capable of pleiotropic effect on all existing color and pattern elements at multiple loci. It should therefore be considered a strong candidate for further studies involving “relationships between spectral and ultrastructure characteristics” in orange ornamentation, and extending to color and/or pattern as a whole as suggested by Kottler (2014). A mechanism is identified by which Pb is capable of balancing overall color and pattern polymorphisms, in turn providing fitness through heterozygosity in diverse complex habitats. We hope that Purple will be mapped to its linkage group.

Throughout this study a recurring thought has consistently arisen in the minds of the author(s). That being, “Where any prior published results based on statistical analysis of quantified male ornaments skewed by failure to identify these two distinct sympatric populations?” In some instances, it would seem XY-linked heritability, brightness, intensity of orange chroma and hue in male ornaments might in an indirect fashion balance out end results. While in others, failure to identify the Pb gene may have biased their results.

## Photo Imaging

Photos by author(s) were taken with a Fujifilm FinePix HS25EXR; settings Macro, AF: center, Auto Focus: continuous, varying Exposure Compensation, Image Size 16:9, Image Quality: Fine, ISO: 200, Film Simulation: Astia/Soft, White Balance: 0, Tone: STD, Dynamic Range: 200, Sharpness: STD, Noise Reduction: High, Intelligent Sharpness: On. Lens: Fujinon 30x Optical Zoom. Flash: External mounted EF-42 Slave Flash; settings at EV: 0.0, 35mm, PR1/1, Flash: -2/3. Photos cropped or brightness adjusted when needed with Microsoft Office 2010 Picture Manager and Adobe Photoshop CS5. All photos by author(s), unless otherwise noted.

## Microscopy

All Digital Image processing by conventional bright and dark field equipment. AmScope M158C. Camera(s): 1. MD35, Resolution: 0.3MP 2. MD200, Resolution: 2MP USB Digital, Sensor: Aptina (Color), Sensor Type: CMOS. Software: AmScope for Windows. An attempt was made to restrict ambient light during both daytime and nighttime imaging of specimens. Imaging was performed with reflected or transmitted practical light sources as indicated. Where delineation in results warranted, a series of three photos from each location were taken and presented in the results; reflected (top light only), transmitted (bottom light only), combined reflected + transmitted (top and bottom light).

For purposes of this study low resolution photos were often preferred over higher resolution for clarity at settings of 40X, 100X or 400X. No images were stained. As identified, individual images are full body (non-dissected), or manually de-fleshed (dissected) skin samples. Samples were air dried for minimal time periods of less than one hour for aid in dissection. All samples and images from right side of body, unless otherwise noted. No cover glass was utilized, to reduce damage to chromatophore shape, structure and positioning. No preservatives were used during imaging, though rehydration was done as needed for clarity. All photos were by the senior author, unless otherwise noted.

## Ethics Statement

This study adhered to established ethical practices under AVMA Guidelines for the Euthanasia of Animals: 2013 Edition, S6.2.2 Physical Methods (6).

All euthanized specimens were photographed immediately, or as soon as possible, after temperature reduction (rapid chilling) in water (H_2_0) at temperatures just above freezing (0°C) to avoid potential damage to tissue and chromatophores, while preserving maximum expression of motile xantho-erythrophores in Pb and non-Pb specimens. All anesthetized specimens were photographed immediately after short-term immersion in a mixture of 50% aged tank water (H_2_0) and 50% carbonated water (H_2_CO_3_).

All dried specimens photographed immediately after rehydration in cold water (H_2_0). Prior euthanasia was by cold water (H_2_0) immersion at temperatures just above freezing (0 °C). MS-222 (Tricaine methanesulfonate) was not used to avoid the potential for reported damage and/or alterations to chromatophores, in particular melanophores, prior to slide preparation.

## Competing Interests and Funding

The authors declare that they have no competing interests. Senior author is a member of the Editorial Board for Poeciliid Research; International Journal of the Bioflux Society, and requested non-affiliated independent peer review volunteers.

The authors received no funding for this work.

## Notes

This publication is number one (1) of four (4) by Bias and Squire in the study of Purple Body (*Pb*) in *Poecilia reticulata*:

1. The Cellular Expression and Genetics of an Established Polymorphism in *Poecilia reticulata*; “Purple Body, (*Pb)*” is an Autosomal Dominant Gene,

2. The Cellular Expression and Genetics of Purple Body (*Pb*) in *Poecilia reticulata*, and its Interactions with Asian Blau (*Ab*) and Blond (*bb*) under Reflected and Transmitted Light,

3. The Cellular Expression and Genetics of Purple Body (*Pb*) in the Ocular Media of the Guppy *Poecilia reticulata*,

4. The Phenotypic Expression of Purple Body (*Pb*) in Domestic Guppy Strains of *Poecilia reticulata*.

## Acknowledgements

To my best friend and wife Deana Bias, for her support and persistence over the last several years in this four part study… To my co-author and dear friend Rick Squire for his patience as a mentor… To those Domestic Breeders who willingly provided additionally needed pedigree strains and study populations for completion of this paper…

## Supporting Information

S1 TABLE 1 – Condensed Breeding’s and Results; male x female test breeding’s of *Poecilia reticulata*. Generations 1,2,3. Charting and results only. (Included as appendix and referenced in text body).

S2 TABLE 2 – Expanded Breeding’s and Results; male x female test breeding’s of *Poecilia reticulata*. Charting, results and photos of all fish. Generations 1,2,3. (Included as appendix and referenced in text body).

S3 - Materials Full Description and Sources.

